# Transcriptome analysis of EPO and GFP HEK293 Cell-lines Reveal Shifts in Energy and ER Capacity Support Improved Erythropoietin Production in HEK293F Cells

**DOI:** 10.1101/2020.09.16.299966

**Authors:** Rasool Saghaleyni, Magdalena Malm, Jan Zrimec, Ronia Razavi, Num Wistbacka, Veronique Chotteau, Diane Hatton, Luigi Grassi, Aleksej Zelezniak, Thomas Svensson, Jens Nielsen, Jonathan L. Robinson, Johan Rockberg

**Author notes:** Corresponding authors: Johan Rockberg, KTH - Royal Institute of Technology, School of Engineering Sciences in Chemistry, Biotechnology and Health, S-106 91 Stockholm, Sweden, Jonathan L. Robinson, Department of Biology and Biological Engineering, Chalmers University of Technology, 41296 Gothenburg, Sweden.

## Abstract

Higher eukaryotic cell lines like HEK293 are the preferred hosts for production of therapeutic proteins requiring human post translational processing. However, recombinant protein production can result in severe stress on the cellular machinery, resulting in limited titre and product quality. To investigate the cellular and metabolic characteristics associated with these limitations, we compared erythropoietin (secretory) and GFP (non-secretory) protein producer HEK293 cell-lines using transcriptomics analysis. Despite the high demand for ATP in all protein producer clones, a significantly higher capacity for ATP production was observed with erythropoietin producers as evidenced by the enrichment of upregulated genes in the oxidative phosphorylation pathway. In addition, ribosomal genes exhibited specific patterns of expression depending on the recombinant protein and the production rate. In a clone displaying a dramatically increased erythropoietin secretion, we detected higher ER stress, including upregulation of the ATF6B gene. Our results are significant in recognizing key pathways for recombinant protein production and identifying potential target genes for further development of secretory power in mammalian cell factories.

**In Brief:** Although the protein secretion process has been widely studied, the complexity of it leaves many questions with regards to defining bottlenecks for successful protein secretion to be answered. By investigating the transcriptomic profiles of different HEK293 clones with varying translational rates producing either the secreted protein erythropoietin or the intracellular GFP, we reveal that high ATP production and improved capacity of specific post-translational pathways are key factors associated with boosting erythropoietin production.

**Highlights:**

- Transcriptomics analysis of a panel of HEK293 stable cell lines expressing GFP or erythropoietin (EPO) at varying translational rates
- Expression of mitochondrial ribosomal genes is positively correlated with EPO secretion
- Expression of different cytosolic ribosomal genes are correlated with productivity in a recombinant-protein specific manner
- High EPO producing clones have significant upregulation of ATF6B, potentially enabling a beneficial ER stress response to cope with high protein secretion

**Graphical Abstract:** 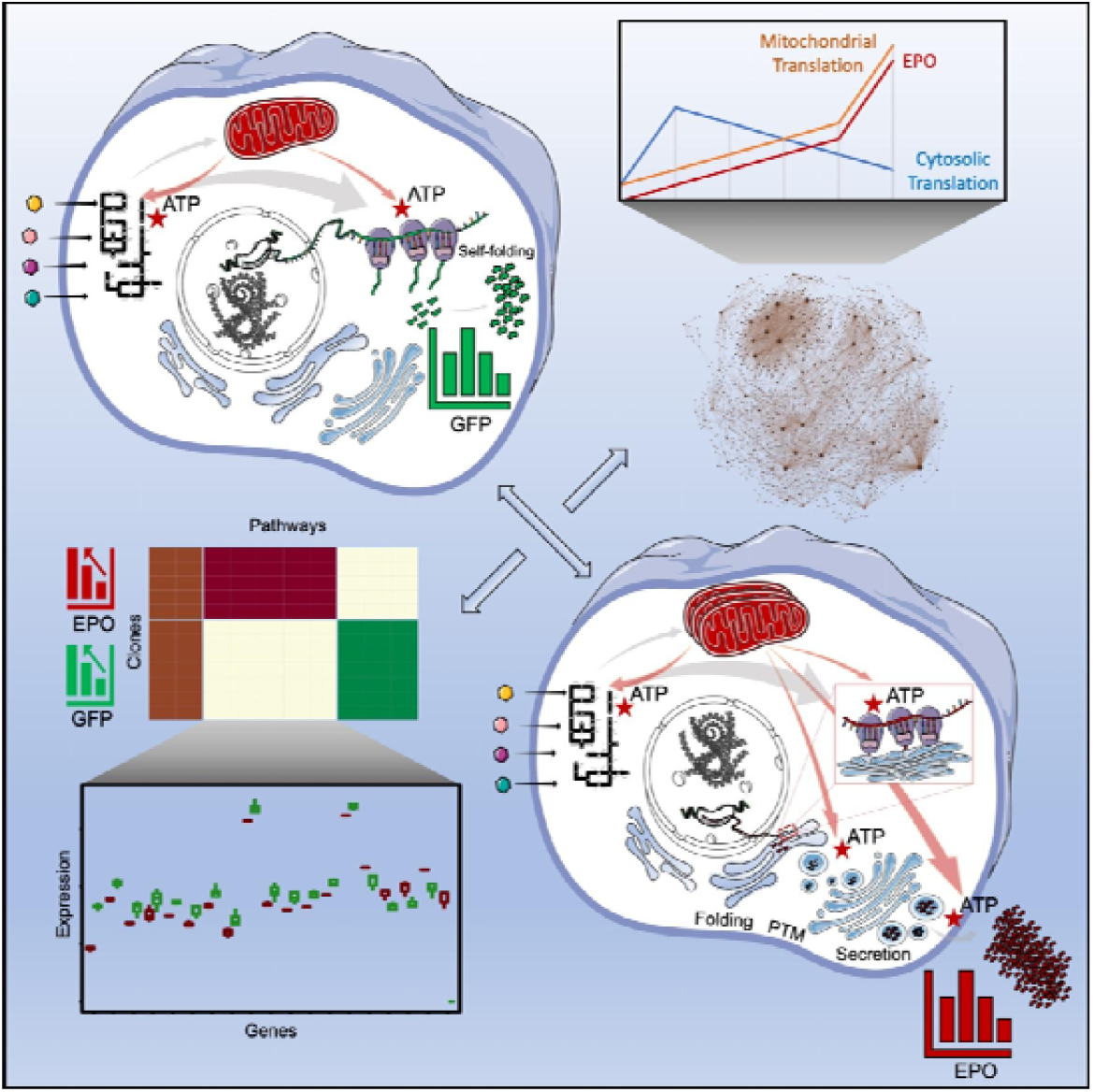

## 1. Introduction

The demand for greater efficiency and quality of protein production in biotechnology is rapidly increasing due to substantial advances in drug discovery (Tambuyzer et al., 2020) and the need for highly effective pharmaceutical proteins for the treatment of severe diseases, such as cancer (Kintzing et al., 2016). Chinese hamster ovary (CHO) cells are the current standard host for the production of a wide range of recombinant proteins partly due to the ability to generate similar post-translational modifications (PTMs) as those in humans, which is often a requirement for complex therapeutic proteins (Davy et al., 2017; Orellana et al., 2015; 2017). However, the PTM pattern of CHO cells is not identical to human PTMs (Dumont et al., 2016) and the incompatibility with some types of proteins negatively affects drug efficacy, potency or stability (Goh and Ng, 2017; Kuriakose et al., 2016). Therefore, besides further development of the CHO cell-line to meet the high-quality PTM requirements and increased yields (Datta et al., 2013; Fouladiha et al., 2020; Koffas et al., 2018; Liang et al., 2020; Tejwani et al., 2018; Wang et al., 2020), a lot of focus on improving hosts for biopharmaceutical production is on cell factories derived from human cells with the natural ability of generating human PTMs, such as the Human Embryonic Kidney 293 (HEK293) cells (Almo and Love, 2014; Malm et al., 2020; Tegel et al., 2020).

Although human derived cell-lines benefit from the ability to generate human PTMs, challenges still remain, such as increasing the protein production titer (Chin et al., 2019; Dietmair et al., 2012; Mori et al., 2020) and creating a genetic engineering toolbox with specialized tools for human cells (Xu and Qi, 2019). Recent publications have pursued some of these challenges, including the aim to increase the protein production and secretion power either by cell-line development approaches (Chin et al., 2019; Rahimpour et al., 2013) or cell culture process optimization (Schwarz et al., 2019), as well as to increase the quality of the secreted proteins by engineering folding and PTM pathways (Behrouz et al., 2020; Del Val et al., 2016; Liang et al., 2020; Meuris et al., 2014). However, despite the current knowledge of protein production and secretion in mammalian cells, there is still notable ambiguity in understanding and predicting the production and secretion rates and product quality under different conditions. This is due to the complexity and presence of many specific biochemical steps across multiple cell organelles that orchestrate, as well as define the rates of production and secretion of each protein (Kaufman and Popolo, 2018). Accordingly, the limited understanding of the biology behind the protein production process, combined with the continuously increasing demands on production quantity from industry and product quality from regulatory bodies, result in a high risk of production failure for many therapeutic proteins.

In the present study, we conducted a transcriptomic comparative analysis to capture physiological differences caused by protein production and secretion in HEK293F cells. In order to understand which differences are caused by protein production and which arise from the secretion-related processes, we generated two groups of cells producing either the secretory protein erythropoietin (EPO) or the non-secretory protein GFP and compared each of these groups with each other and with their parental cell-lines. We found genes with a high correlation with EPO or GFP production and investigated biological functions associated with them. Furthermore, we investigated ribosome heterogeneity between EPO and GFP producers and detected ribosomal genes with specific patterns of expression correlating with EPO and GFP production. Since the generation of single clones by random integration enabled us to isolate a clone with greatly increased EPO production titer - a 3-fold increase compared with the other clones - we set out to identify the reasons behind these improved protein titers, highlighting genes that can potentially facilitate increased protein production and secretion in future studies.

## 2. Results

### 2.1 Single cell cloning generates EPO and GFP producer clones with an altered metabolism from the host cell-line

To investigate recombinant protein production in HEK293 cells, we transformed 293-F cell-lines to generate secretory protein (EPO) producer clones and non-secretory protein (GFP) producer clones, respectively (**Figure 1A, M1**-**3**). Initially, polyclonal batches of cells producing either EPO or GFP (EPOpoly and GFPpoly, respectively) were generated through random integration of plasmid DNA into the host genome resulting in collections of clones with various transgene integration sites and copy numbers. From these batches five EPO-producing and seven GFP-producing clones were isolated. We observed that the growth rates of the EPO and GFP producers were lower than those of their respective host cell lines, but the decrease was no more than 22% and 14% compared to the hosts, respectively (**Figure 1B**). Protein productivity was however markedly different among clones and varied by almost 6-fold with EPO producers and up to 4-fold with GFP producers (**Figure 1C-D, M2**). The most productive EPO clone (EPOF21) had a titer of 13.9 pg/cell.day, over 3-fold higher than the second highest producing clone EPO8 (productivity = 4.05 pg/cell.day) and, interestingly its mRNA copy number was up to 20% lower (**Figure 1E, Figure S1A, M4**). Except for this extraordinary production clone, we could measure significant correlation (Pearson’s *r* = 0.99, *p* = 0.001) between EPO mRNA amount and secreted EPO productivity for all clones (**Figure 1E, Figure S1B**), which was similarly high as the correlation observed between mRNA copy number and GFP productivity (Pearson’s *r* = 0.88, *p* = 0.004, **Figure S1C**). For both EPO and GFP clones, no significant correlation was found between protein productivity and gene copy number or clone growth rate (**Figure S1A-C**).

**Figure 1.**
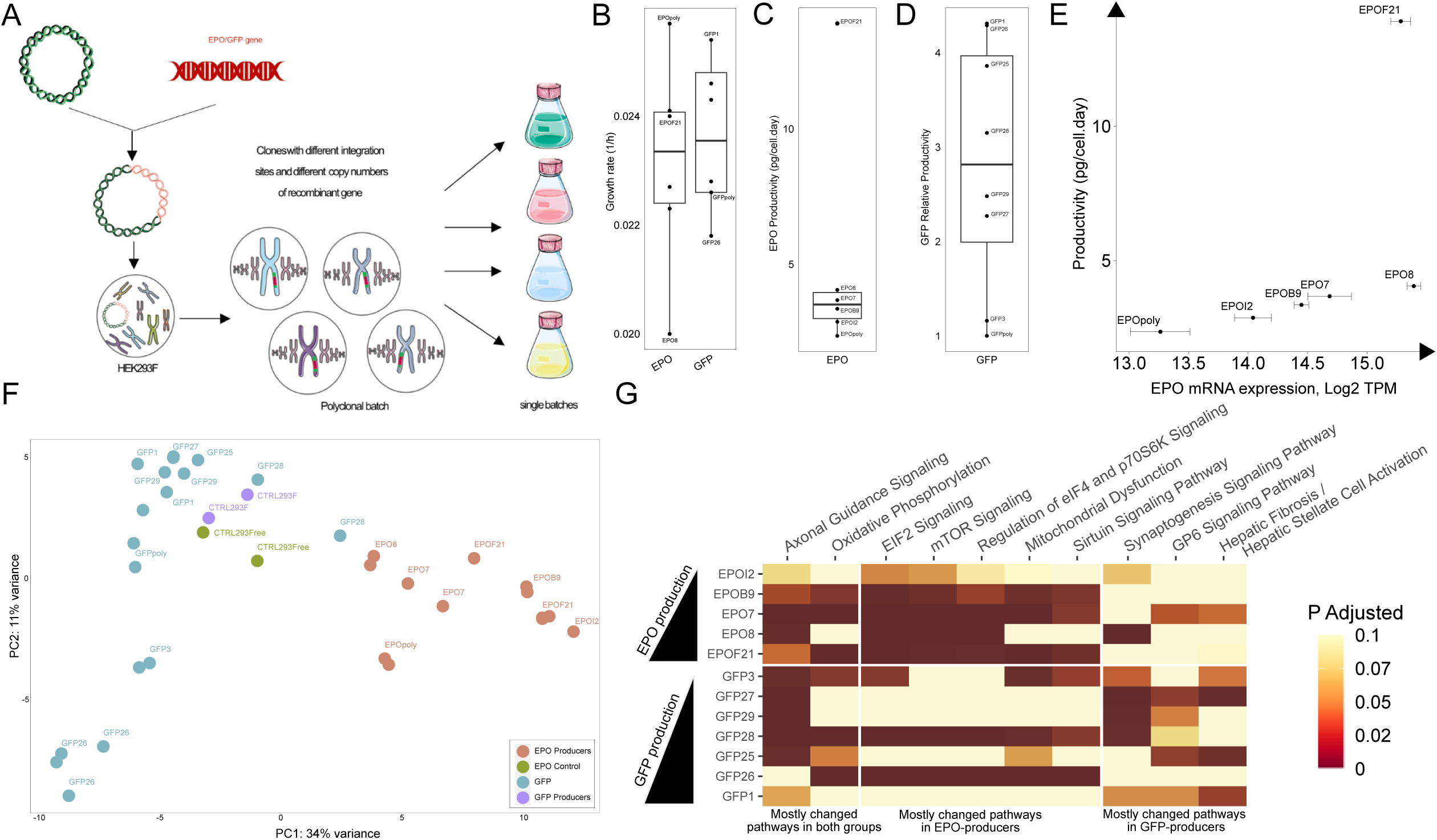
Single cell cloning procedure generates EPO and GFP producer clones with an altered metabolism from the host cell-line. (A) Schematic diagram of the cell line development resulting in random integration sites and copy number of the EPO and GFP genes across the genomes. (B) Growth rates of EPO and GFP producer clones. (C) Specific productivity of EPO in different HEK293 clones. (D) Relative productivity of GFP in the different clones in comparison to the polyclonal batch of the GFP clones (GFPpoly). (E) Specific EPO productivity versus its mRNA expression in different clones. (F) Principal component one in PCA analysis separates producer clones based on their recombinant EPO or GFP protein. (G) Most significantly (B.H. adj. *p*-value < 0.05) enriched pathways in pairwise comparison of EPO or GFP producers against the control hosts.

We analysed and compared transcriptomic data (Illumina HiSeq, **M1**) to find how the protein producer cell lines differ from their respective parental host cell lines, **Table S1**). Principal component analysis (PCA) clustered clones in the first component based on their respective recombinant protein (EPO or GFP, **Figure 1F, Figure S1D**). We performed pairwise differential expression analysis between each recombinant protein producer clone and their respective host (**Figure S1E-G, M5-6, Table S2**). Results of this analysis (**Figure 1E-F**) showed, while in EPO producers EPOI2 had the highest number of differentially expressed (B.H. adj. *p*-value < 0.05, |L2FC| > 1) genes (1137 genes upregulated and 82 genes down regulated), in GFP producers GFP26 clone was the most different clone compared to control cell-line (487 genes down regulated, 466 genes upregulated) based on the number of differentially expressed genes. We also found 45 and 10 common differentially expressed genes in pairwise comparison of EPO producers and GFP producers with their respective control cell line, respectively (**Figure S1G)**.

Pathway enrichment analysis (**Figure 1G**) showed that axonal guidance signaling and oxidative phosphorylation were the two pathways in the IPA pathways database exhibiting constant change by most of the recombinant protein producer clones. Apart from these two pathways, eIF2 signaling and mTOR signaling pathways were significantly (B.H. adj. *p*-value < 0.05) altered across all EPO producers (**Figure 1G**). Moreover, mitochondrial dysfunction, regulation of eIF4, p7056K signaling and Sirtuin signaling pathways were significantly (B.H. adj. *p*-value < 0.05) altered in more than half of the EPO producers. On the other hand, gene enrichment analysis of GFP producers against their parental HEK293 cell-line did not reveal any common pathways enriched across all the cell lines (**M6**). Instead, we observed generally different phenotypes between clones, with the exception of axonal guidance signaling, oxidative phosphorylation, synaptogenesis signaling and GP6 signaling pathways with a significant change (B.H. adj. *p*-value < 0.05) in more than half of the GFP producers (**Figure 1G**).

To further investigate the direction of change and the downstream effects of gene expression changes in altered signaling pathways, we performed Ingenuity Pathway Analysis (IPA, QIAGEN Inc.) (Krämer et al., 2014), to infer downstream links of gene expression effects. IPA indicated that significantly upregulated (B.H. adj. *p*-value < 0.05) genes related to translational and post-translational pathways were enriched in eIF2 and mTOR signalling pathways in EPO producers compared to the control cell line (**Figure 1G, Figure S2, M6**). An increase in the expression of genes associated with translation was predicted to activate downstream processes including protein folding, ER stress and apoptosis as well as upstream processes, such as amino acid biosynthesis (**Figure S2**).

### 2.2 Expression of associated genes with mitochondrial ribosomal proteins is positively correlated with EPO production

We sought to find genes that significantly correlate with EPO and GFP production and investigate their roles in the process of protein production (**M4**). For this purpose, we first extracted genes with an average TPM above 10 across all cell lines for both EPO and GFP and then considered positively and negatively correlated (|Pearson’s *r*| > 0.5, *p* < 0.05) genes. Altogether, 223 and 93 genes were positively and negatively correlated with EPO production, respectively, and 99 and 19 genes were positively and negatively correlated with GFP production, respectively (*Mean*_*TPM*_ > 10, |*r*| > 0.5, *p* < 0.05) (**Figure 2A, Table S3**). Amongst these genes, 6 genes were found to correlate (|*r*| > 0.5, *p* < 0.05) with both EPO and GFP (**Figure 2B-C**). Their known functions suggested that they might generally be involved in regulation of protein production; the function of MVP is related to generating ribonucleoproteins (Zheng et al., 2005), B3GNT5 is involved in posttranslational modifications (Togayachi et al., 2001), and for the less characterized gene products of WDR53 and NPAS1, evidence suggests that they act as regulatory elements in cell metabolism (Wu et al., 2016). Analysis of the corresponding biological GO terms (HyperGSA, *p* < 0.05, **M4**) for the EPO-correlating genes showed that many of the genes with a positive correlation (*r* > 0.5) are involved in secretory pathway components such as protein folding, post-translational protein modification and post-Golgi mediated transport (**Figure 2D, M4**). Moreover, a major node enriched with both negative and positive EPO-correlating genes was associated with mitochondrion organization. Conversely, the GO terms (HyperGSA, *p* < 0.05, **M4**) associated with GFP-correlating genes spanned genes that were mostly negatively correlated (*r* < −0.5) with GFP production and covered a wide range of GO terms including RNA catabolic processes, RNA export from the nucleus and translation (**Figure 2E)**. One of the detected enriched pathways by both positive and negative GFP-correlating genes was translation (HyperGSA *p*-value *=* 0.029) and interestingly, almost half of the genes in this pathway (4 out of 9) were components of the large subunit of cytosolic ribosomes (RPL3, RPL4, RPL29, RPL36A).

**Figure 2.**
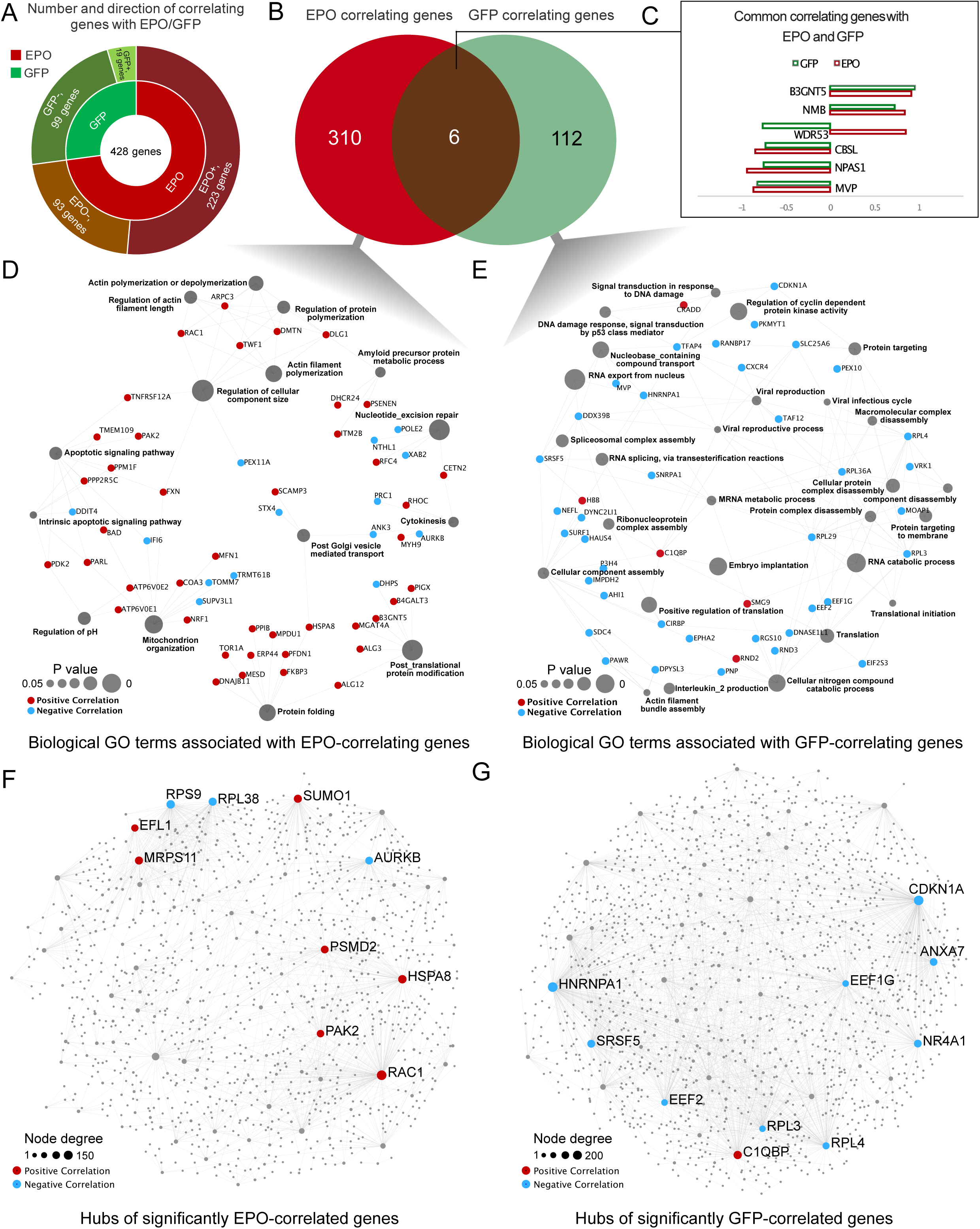
Translational and post-translational genes are strongly correlated with EPO and GFP production. (A) Most significantly (*Mean*_*TPM*_ > 10, |Pearson’s *r*| > 0.5, *p*-value < 0.05) correlated genes with EPO or GFP production. Positive and negative signs show positive and negative correlation, respectively (B) Common and specific correlated genes with EPO and GFP production (C) Correlation coefficient of common correlated genes with both EPO and GFP production. (D-E) Enrichment analysis of significantly correlated (*Mean*_*TPM*_ > 10, |*r*| > 0.5, *p* < 0.05) genes with EPO (D) and GFP production (E). (F-G) Top ten hubs with the highest number of interactions (supported by experimental evidence outlined in STRING database) among significantly correlated (*Mean*_*TPM*_ > 10, |*r*| > 0.5, *p* < 0.05) genes with EPO (F) and GFP (G) productivity.

To find the major regulators among the genes highly correlated (*Mean*_TPM_ > 10, |*r*| > 0.5, *p* < 0.05) with either EPO or GFP productivity, we generated interaction networks between the genes and their first-order interacting partners based on experimental evidence (confidence score > 900) extracted from the STRING database (Szklarczyk et al., 2019) (**Figure 2F-G, Table S3, M4**). We excluded those interacting genes that are not expressed in our dataset or have very low expression (*Mean*_TPM_ < 10) and then ranked the genes based on their node degree (*k*), which measures the interactivity of each gene based on the number of observed interacting gene partners. The node degree (*k*) of the top ten most interactive genes (**Figure 2F-G**: network hubs) ranged from 53 to 144 and 79 to 202 in the networks of EPO- and GFP-correlating genes, respectively (**Table S3)**. Among the top ten interacting genes in the network of EPO-correlating genes, four of them were directly involved in translation. RPL38 (*r* = −0.89, *p* = 0.01, *k* = 73) and RPS9 (*r* = −0.91, *p* = 0.01, *k* = 88), both components of cytosolic ribosomal subunits, were negatively correlated with EPO production (Kondrashov et al., 2011), while MRPS11 (*r* = 0.9, *p* = 0.01, *k* = 71), a mitochondrial ribosomal gene, and EFL1 (*p* = 0.82, *r* = 0.04, *k* = 48), involved in 60S ribosomal subunit biogenesis (Thomson et al., 2013), were positively correlated. Similarly, with GFP producers, we observed a negative trend of expression in genes involved in translation with increasing GFP production. Likewise, four out of the top ten genes (EEF2, EEF1G, RPL3 and RPL4) with the highest number of interactions in the network of GFP-correlating genes, serving as translation factors or components of the ribosomal large subunit, were negatively correlated (*r* < −0.7, *p* ≤ 0.05, 94 < *k* < 202) with GFP production. When looking into all the ribosomal genes correlating (|*r*| > 0.5, *p* < 0.05) with EPO production **(Figure S3, M4**), besides RPL38, RPS9 and MRPS11 mentioned above, there were additionally three ribosomal genes (MRPS18A, MRPL40 and MRPL49) belonging to mitochondrial ribosomal genes positively correlating (r > 0.8, *p* < 0.05) with EPO production (**Figure S3A)**. Similarly in GFP producers, beside RPL3 and RPL4, three ribosomal genes (EEF2, RPL29 and RPL36A, all components of cytosolic ribosome), were significantly negatively correlated (*r* < −0.7, *p* ≤ 0.04) with GFP production **(Figure S3B**).

Among the other (non-ribosomal) top ten interacting genes in the network of EPO-correlating (|*r*| > 0.5, *p* < 0.05) genes, RAC1 (*r* = 0.84, *p* = 0.03, *k* = 144) is a member of the rho family of GTPases and function as molecular switches that regulate a variety of different processes within the cell, including extracellular organization and cell division (Lin et al., 2005; Thomas et al., 2019). PAK2 (*r* = 0.84, *p* = 0.03, *k* = 53), one of the other genes with a high and positive correlation with EPO production, is a direct target of RAC1 and links Rho GTPases to cytoskeleton reorganization and nuclear signaling (May et al., 2014). Furthermore, we observed positive correlation in EPO production and expression of AURKB (*r* = 0.84, *p* = 0.03, *k* = 62), a serine/threonine kinase, which with RAC1 and PAK2 is active in the regulation of chromosomes segregation during mitosis and meiosis through association with microtubules (May et al., 2014). The wide range of targets for these three genes (RAC1, PAK2 and AURKB) and positive correlation of their expression with EPO production (with exactly the same correlation coefficients for all of them) suggests coordinated expression of these genes could be related to EPO production process. We also detected positive correlation of HSPA8 expression with EPO production (*r* = 0.84, *p* = 0.03, *k* = 84). HSPA8 is a well-known chaperone acting in the protein folding process and previously shown to increase protein production by improving the folding processes in CHO cells (Bonam et al., 2019; Lee et al., 2009). In the network of GFP correlated genes we SRSF5 (*r* = −0.71, *p =* 0.04, *k* = 87) and HNRNPA1 (*r* = −0.7, *p =* 0.04, *k* = 83), which are involved in the metabolism of pre-mRNA both shown negative correlation with EPO production (Chen et al., 2018; Yang et al., 2018). This suggests that a reverse trend in expression of translation-associated genes with GFP production could be coupled with upstream pathways involved in mRNA maturation.

### 2.3 Erythropoietin production requires restructuring of cellular metabolism to meet its energy demands

We sought to understand which changes in protein producers were due to the production of secretory EPO and which were common to all hosts. Principal component analysis of the EPO and GFP transcriptomics data (**M5**) marked differences between EPO and GFP producers (**Figure 1F**), where a complete separation of EPO and GFP clones with the 1st principal component (34% of variance explained) showed that the transcriptomics data could capture differences between the secretory EPO and non-secretory GFP producers. We found 986 (922 up- and 64 down regulated) differentially expressed genes (adj *p*-value < 0.05, |Log2 Fold Change| > 1) between EPO and GFP producers. Gene set enrichment analysis (**Figure S4A, M7**) showed that with EPO producers, beside gene sets specific for secretory protein production, such as proteins targeting the endoplasmic reticulum or the cell membrane, the oxidative phosphorylation pathway was significantly (B.H. adj. *p*-value < 0.05) enriched by up regulated genes (**Figure 3A**).

**Figure 3.**
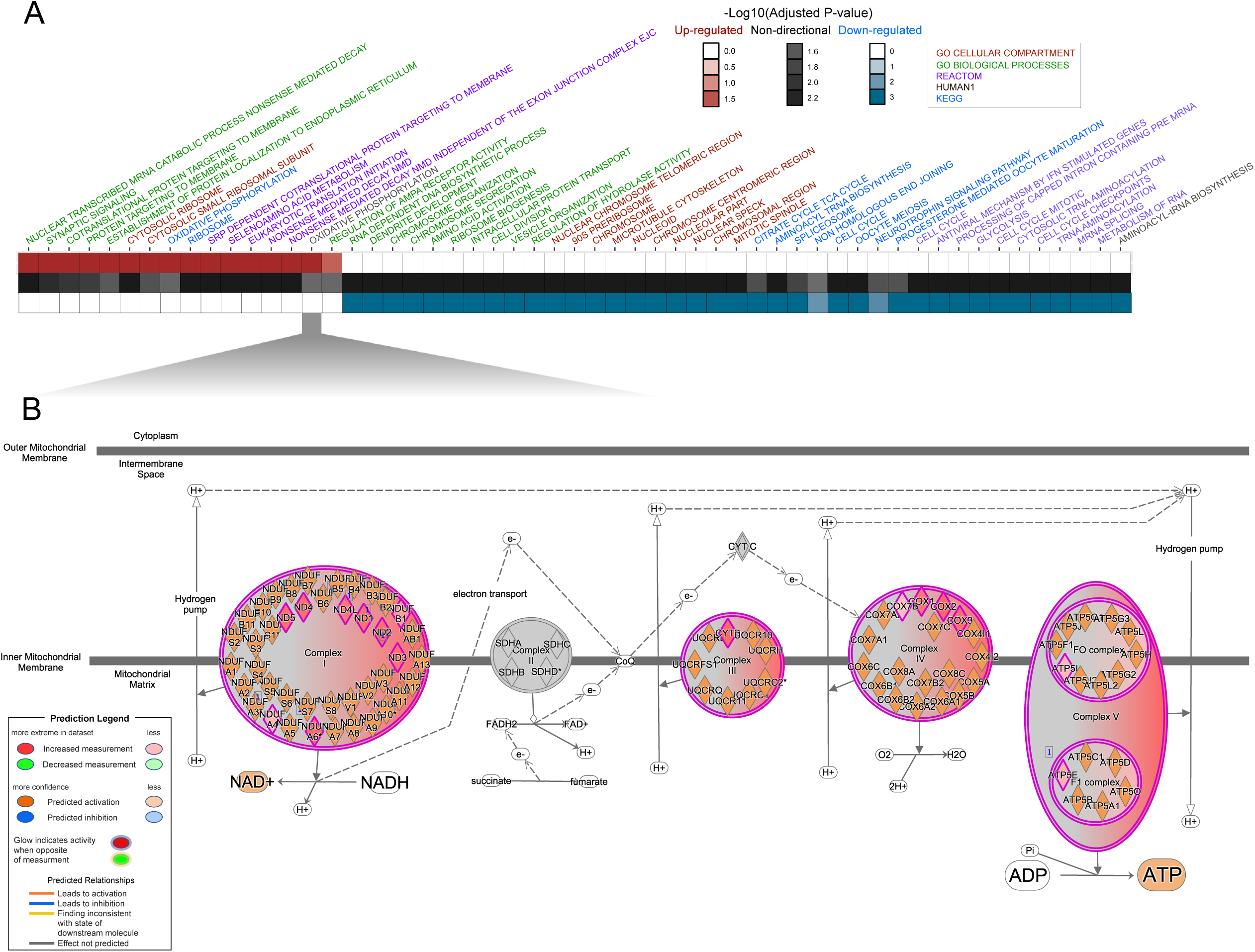
Cellular metabolism is restructured to meet the energy demands of EPO production. (A) Heatmap of enriched (B.H. adj. *p*-value < 0.05) pathways in comparison of EPO producers with GFP producers. Red and blue color ranges show up- and down regulation in EPO producers, respectively. Also, column color shows corresponding gene set collection for enriched pathway (B) The oxidative phosphorylation pathway was significantly (B.H. adj. *p*-value = 0.03) up regulated in EPO producers compared to GFP producers. Detailed visualization of IPA prediction indicates up regulated genes and their role in increasing ATP production.

To further investigate the increased oxidative phosphorylation activity among EPO producers, we used IPA to elucidate how expression changes in genes associated with this pathway impart their effect and which mitochondrial membrane complexes are more active in increasing ATP production (**Figure 3B**). Indeed, all differentially expressed genes (B.H. adj. *p*-value < 0.05) involved in oxidative phosphorylation exhibited increased expression in EPO producers. The up regulated genes are associated with all complexes in the electron transport chain except for complex II, which facilitates the donation of electrons from FADH2. IPA clearly indicated higher activity in NADH dehydrogenase (complex I) to create an electrochemical gradient across the inner mitochondrial membrane followed by an increased activity of complex III to complex V and finally higher ATP production. All genes of mitochondrial origin across the electron transport chain (ETC) complexes exhibited at least a 2-fold increase in their expression (**Figure S4B**: genes with names starting with ‘MT-’). Apart from up regulation of mitochondrial genes, other genes that are expressed from the nuclear genome and are associated with oxidative phosphorylation also had a significant (B.H. adj. *p*-value < 0.05) expression increase in EPO producers.

Moreover, pathways related to translation and ribosome biogenesis were among the most significantly enriched (B.H. adj. *p*-value < 0.05) by upregulated genes in both groups of EPO and GFP producers (**Figure 3A**). This suggested that, in general, gene expression associated with translation processes is adopted in each group of EPO and GFP producers to support their specific needs imparted by their particular recombinant protein. To follow this observation, we investigated the expression of ribosomal genes in pairwise comparison of each clone with its respective control (**Figure S5A-B**). Although GFP producers did not share common differentially expressed ribosomal genes (B.H. adj. *p*-value < 0.05, **Figure S5C**), EPO producers showed 22 common differentially expressed (*p*-value < 0.05) ribosomal genes with at least 50% increase in their expression in comparison to control, **Figure S5D**). Functionality of these genes is mostly related to SRP-dependent co-translational protein targeting to membrane. This suggests EPO-producers have increased the share of genes related to co-translational protein targeting to membrane in their ribosomes to facilitate EPO production.

### 2.4 Post-translational pathways can limit the EPO protein production rate

Besides the observations regarding differences between mRNA copy number and protein productivity in EPOF21 in comparison to other EPO producers (**Figure 1E**), differences in transcription and translation as well as post-translational pathways might have affected protein productivity. In order to find genes and pathways with altered patterns of expression, we first conducted pairwise differential expression analysis to find which genes are differentially expressed between EPOF21 and other EPO producers (**Figure S6A, M5**). Results of differential expression analysis indicated EPO expression is not significantly changing between clones. Number of down regulated differentially expressed genes (B.H. adj. *p*-value < 0.05, L2FC < −1) varied between 186 (EPOF21 vs. EPOI2) and 491 (EPOF21 vs. EPO8) and number of up regulated genes (B.H. adj. *p*-value < 0.05, L2FC > 1) showed a variation between 113 (EPOF21 vs. EPOI2) and 792 (EPOF21 vs. EPO8) genes (**Figure S6A)**. We also found 44 commonly differentially expressed genes in comparisons of each of EPO producer clones against EPOF21. Gene set enrichment analysis (**M7**) between EPOF21 and other EPO producer cell-lines (**Figure S6B-C**), indicated many common changes between EPOF21 and the other EPO producers were related to post-translational pathways. To obtain an extended gene set spanning pathways related to protein secretion, we used a set of secretory protein machinery genes that are defined as core genes involved in protein secretion in human cells (Feizi et al., 2017; Gutierrez et al., 2020), and then searched for all GO pathways (Liberzon et al., 2011) that contained a significant (B.H. adj. *p*-value < 0.05) number of these genes (HyperGSA, **M8**). This collection of gene sets was then used to compare the different EPO producer cell-lines (**Figure 4A**: top altered pathways in at least one of the comparisons). Interestingly, when comparing EPOF21 to all other cell lines we observed marked differences between the EPO8 cell-line and the other EPO producers, which could have resulted from differences in EPO mRNA expression levels between EPO8 and the other cell-lines (**Figure 1E**). Furthermore, comparison of the up regulated pathways in EPOF21 against other EPO producers except EPO8 indicated a marked expression increase in gene sets related to the endoplasmic reticulum and handling of misfolded proteins in EPOF21 (**Figure 4A**). Although we did not observe any significantly down regulated pathways in EPOF21 in comparison to EPOB9 and EPOI2, some pathways related to protein localization to organelles and also cellular macromolecular catabolic processes were down regulated in EPOF21 in comparison to EPO8 and EPO7. This may indicate the potential problematic steps in the process of folding and targeting of secretory proteins in EPO8 and EPO7 that result in the lower EPO production rates in these cell-lines relative to EPOF21.

**Figure 4.**
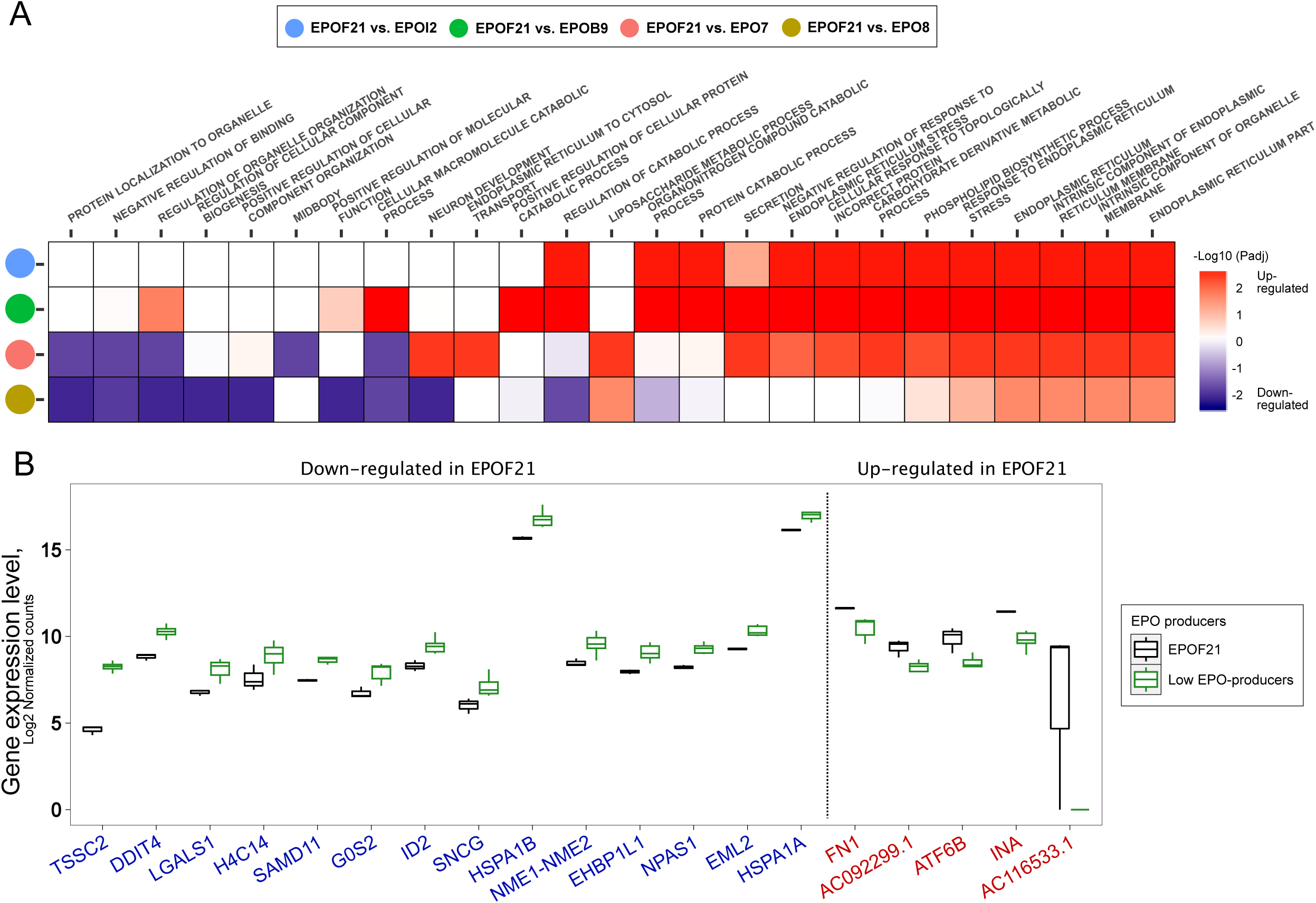
Post-translational pathways are different between EPO producer clones. (A) Significantly enriched GO terms in pairwise comparisons of EPOF21 and other EPO producers using GO slim secretion. (B) Gene expression levels of the most significant (B.H. adj. *p*-value < 0.05) consistently differentially expressed genes between EPOF21 and other EPO producers.

To further investigate specific differences of EPOF21 with other EPO producers, we analysed which differentially expressed genes exhibited a significant and solid pattern of change (B.H adj. *p*-value < 0.05, L2FC > 1, *Mean*_*TPM*_ > 10). Of the 19 genes that displayed a consistent pattern of change, 5 were up regulated and 14 down regulated (**Figure 4B**). Among the upregulated genes, ATF6B is a transcription factor active under ER stress conditions due to accumulation of unfolded proteins (Thuerauf et al., 2004), whereas FN1 and INA have a role in extracellular matrix assembly (Singh et al., 2010). The gene with the highest positive fold-change between EPOF21 and other EPO producers, AC116533.1 is a pseudogene of the ribosomal gene RPL36A, a ribosomal protein shown to play a role in ribosome biogenesis in yeast (Wan et al., 2015). Down regulated genes covered a wider spectrum of pathways including: apoptosis and growth regulation (LGALS1, G0S2 and EML2), nucleosome organization (HIST2H4A and SAMD1), metabolism of nucleotides (NME1-NME2), protein folding (HSPA1A and HSPA1B) and vesicle trafficking (EHBP1L1). Also, ID1 and NPAS1 play roles in transcription regulatory pathways (Erbel-Sieler et al., 2004; Sikder et al., 2003) and TSSC2 is a pseudogene and a homologue to the Asparagine-Linked Glycosylation 1 (ALG1) gene in yeast (Jaeken et al., 2015).

## 3. Discussion

In the present study, we generated two groups of clones producing either EPO or GFP protein at different levels (**Figure 1A**). EPO and GFP were chosen as well studied model proteins with different characteristics such as final cellular location in order to study the pathways behind protein production and differences caused by protein secretion. Indeed, analysis of the clonal transcriptomics data, where both clonal variation and the recombinant protein type were captured by the most informative first principal component (**Figure 1F**), suggests that the design of the experiment was appropriate for exploring both the protein production and secretion stages. However, despite individual HEK293 clones showing different production titers as well as different gene expression profiles (**Figure 1C-E, Figure S1A**), both transcription and translation were very well synced (**Figure S1B**). Thus, all of the clones except EPOF21, a clone with over 3-fold higher production levels compared to the next highest EPO producer (**Figure 1E)**, had an almost equal ratio of secretion of the recombinant protein versus transcription of the recombinant gene transcripts (**Figure 1E**). The high increase of this ratio in EPOF21 (in relation to the other clones) indicated that major translational and post-translational processes were affected. This suggests that, whereas all clones were useful to study the effects related to non-secretory and secretory protein production as well as their differences, the extraordinary EPOF21 clone is a highly useful candidate to further pinpoint major limiting parameters within secretory protein production, which can in general help to improve the overall productivity of cell factories.

Analysis of transcriptomic data between protein producer hosts and control parental cells showed up regulation of oxidative phosphorylation in all clones regardless of transgene or level of recombinant protein production (**Figure 1G**), which indicates a high energy demand is required to support transgene expression. Moreover, significant upregulation of genes linked to oxidative phosphorylation was observed when comparing EPO producers with GFP producers (**Figure 3B**), suggesting that EPO clones have an even higher energy demand imposed by post-translatory secretory pathways (Gutierrez et al., 2020). Simultaneous up regulation of genes from both the mitochondria and nucleus highlight that the increase in energy production is not governed merely by the change in the mitochondrial genome, but also by the upstream regulatory pathways of the cell (**Figure S4B**). The demand for more energy in EPO producers compared to GFP producers, could be related to the intrinsic molecular differences between EPO and GFP or could be the result of the general absence of secretory energy requirements in GFP producers. For instance, there are 114 different N-linked and O-linked reported structures for EPO (Alocci et al., 2019) but the complexity of post translational processing for GFP is simpler in general (Barondeau et al., 2003).

We note that the EPO protein itself acts as a signaling molecule and could therefore impart regulatory effects on EPO producing clones, such as increased oxidative phosphorylation activity (Plenge et al., 2012). However, EPO activity relies on the presence of the erythropoietin receptor (EpoR) which is generally restricted to erythroid progenitor cells. Evidence supporting protein expression of EpoR and/or EPO activity via EpoR interaction in other (non-hematopoietic) cell types is lacking (Elliott and Sinclair, 2012), with the exception of skeletal muscle (Rundqvist et al., 2009). Furthermore, a study of EpoR expression and activity by Ott et al. (Ott et al., 2015) detected no EpoR protein in HEK293 cells unless they were transfected with an EpoR overexpression vector. In line with this the EPOR expression in our dataset is not correlated with EPO production (**Figure S7A**) and furthermore there is no significant difference in the expression of this gene between EPO producers and GFP producers and also between EPO producers and their control cell-line (**Figure S7A-D**). Moreover, we didn’t detect a solid and significant difference (B.H adj. *p*-value < 0.05, L2FC > 1) in expression of downstream genes in the EPOR signaling pathway (**Figure S7E**) in comparing the expression of these genes between EPOF21 and other EPO producers (**Figure S7F**). We therefore reason that the transcriptional changes observed in EPO-producing clones are likely to be associated primarily with differences in secreted protein production rather than EPO-EpoR signaling, though some confounding effect from signaling cannot be ruled out entirely.

In EPO-clones, significant up regulation of genes belonging to the eIF2 and mTOR signaling pathways could lead to activation of translation (**Figure S2A**). The eIF2 initiation complex is active in translation initiation in eukaryotic cells and plays a role in stabilizing the preinitiation complexes through binding to mRNA, GTP, methionine tRNA and finally the 40S ribosomal subunit, to generate the 43S pre-initiation complex (Hinnebusch, 2011; Schmitt et al., 2010; Stolboushkina and Garber, 2011; Wek et al., 2006). Likewise, activation of mTORC1 module in mTOR signaling pathway (**Figure S2B**) could promote protein synthesis and profoundly increase cellular ATP level by controlling mitochondrial biogenesis (Laplante and Sabatini, 2009). However, when looking into associated GO terms with all positively and negatively correlated genes with recombinant protein production, only GFP production showed a positive correlation with translation-associated genes, whereas EPO production was positively correlated with members of protein folding and post-translational modification pathways (**Figure 2D**). We also observed a positive correlation between expression of genes associated with apoptosis signaling and EPO productivity, which emphasize how protein production pressure could stimulate pathological ER stress and activate apoptotic related pathways (Iurlaro and Muñoz-Pinedo, 2016; Sano and Reed, 2013). Three genes RAC1 and PAK2 detected between the top 10 genes with the highest number of interactions (based on experimental evidence) with other correlating genes (**Figure 2F, M4**), has frequently been reported in previous studies as a module that greatly regulates cytoskeleton reorganization, intracellular trafficking and apoptosis (Chi et al., 2013; Coleman and Olson, 2002; Croisé et al., 2014; Embade et al., 2000). Considering their upstream regulatory roles in controlling cell behaviour and the fact that all these three genes have a significant positive trend of expression with increase in EPO production, it is very likely that upregulation of these genes in high producer cells is associated with their potential role in boosting the level of EPO production.

Among other top ten interacting genes in the network of EPO correlating genes (**Figure 2F**), HSPA8, a member of the heat shock proteins family A, has also been previously reported as a target for improving recombinant protein production in CHO cells (Lee et al., 2009). Likewise, RPL38 and RPS9 (cytosolic ribosomal) and MRPS4 (mitochondrial ribosomal) genes with negative and positive trends of expression with EPO production, respectively, were found among the top ten EPO correlating genes with highest interactions with other genes. Due to similar observations for cytosolic ribosomal genes in GFP producers (**Figure 2G**) and since a negative trend of expression for ribosomal genes with increase in protein production was unexpected, we further investigated this observation by analysing the expression trends for each ribosomal gene that significantly correlated with protein production (**Figure S3**). Remarkably, we detected similar negative and positive trends of expression across all cytosolic and mitochondrial ribosomal genes, respectively. This consistent pattern of change between ribosomal genes belonging to two different compartments indicates the presence of an upstream regulation in favor of a specific need by the cell. The observations that EPO production imposes a high energy burden on the cells (**Figure 1G, Figure 3A-B** and **Figure S5**) and the elevated level of expression of mitochondrial ribosomal genes with higher rates of EPO production and also lack of such observation in GFP producers, suggests that secretory EPO-producing cells allocate more protein resources to produce mitochondrial ribosomes and increase the production of mitochondrial proteins involved in the energy production process through oxidative phosphorylation. These results further highlight the importance of energy metabolism for secreting higher levels of EPO and could explain a general strategy employed by cells to support higher levels of secretory protein production by increasing energy production and compensating for this increase by downregulating the production of cytosolic ribosomes.

We also detected a negative trend in the expression of some cytosolic ribosomal genes (**Figure 2E, Figure 2G** and **Figure S3B**) with increasing GFP production levels. The observed decrease in the expression of some ribosome-associated genes, could be a result of an induced stress due to higher levels of translation, or it could indicate a rearrangement in the profile ribosomal components. The latter is known as ribosome heterogeneity (Genuth and Barna, 2018a) and considers ribosomes as dynamic macromolecular complexes that use a variation of different components in their structure to fit with desired specialized functions (Genuth and Barna, 2018b). It has previously been shown that specialized ribosomes can preferentially translate different subsets of mRNAs (Shi et al., 2017). Our observations highlight the presence of both up regulated and down regulated ribosome-related pathways in EPO and GFP producers (**Figure 3A**) and pinpoint transgene-specific correlations between expression levels of different ribosomal genes and the respective recombinant protein titers (**Figure 2D-E, Figure S3**). These results could suggest that cells decrease the expression of some ribosomal components in a transgene-specific manner, in favor of more convenient production of the specific recombinant protein.

Focusing on the differences of high producer EPOF21 clone with other EPO producers, we observed that the majority of differences among the EPO producer clones are in the post-translational steps, including protein folding, post-translational modifications and handling of misfolded proteins (**Figure 4, Figure S6B-C**). Indeed, in accordance with the increase of EPO transcripts, post-translational and ER-related pathways in EPOF21 were upregulated (**Figure 4A**). However, the EPO8 clone, with almost the same level of EPO transcripts as EPOF21 (**Figure 1E**), did not display an increased activity in some of ER-associated pathways. This indicates that a higher expression of the recombinant gene without the support of post-translational steps is not only inefficient, but can even cause problems with protein expression and lead to lower protein productivity.

To find regulatory elements behind higher activity of post-translational pathways in EPOF21 we investigated genes with a consistent pattern of change between EPOF21 and other EPO-producers. Among the top up regulated genes in EPOF21 was ATF6B (**Figure 4B**). ATF6B is integrated within the ER membrane under normal conditions, but during ER stress conditions, the cytoplasmic N-terminal domain is cleaved and the protein enters the nucleus to activate ER stress response genes (Iurlaro and Muñoz-Pinedo, 2016; Thuerauf et al., 2004). Although ATF6B and its isomer ATF6A both activate the ER stress response genes (ERSRG), ATF6B represses the strong effect of ATF6A, and through this regulation causes a moderated activation of the ER stress response in comparison to ATF6A (Correll et al., 2019; Koul et al., 2017). Moreover, a previous study indicated that targeting ATF6A using a microRNA (miR-1287) enhances productivity in CHO cells producing a therapeutic antibody (Pieper et al., 2017), where miR-1287 had a very similar role to ATF6B in competing with and suppressing ATF6A to decrease the ER stress response. Additionally, it has previously been reported that continued ER stress causes higher expression of genes involved in the folding process (Jäger et al., 2012). Accordingly, activation of ATF6A during unfolded protein response (UPR) condition induces upregulation of heat shock proteins such as HSPA1A and HSPA1B (Gargalovic et al., 2006; Lee et al., 2003). So, downregulation of the stress-inducible chaperones HSPA1A and HSPA1B in EPOF21 (**Figure 4B**) could suggest moderate level of ER stress in the this clone, capable of supporting a high level of protein secretion, potentially governed by the higher expression of ATF6B. So, the difference in the expression levels of ATF6B transcription factor between EPOF21 and the low EPO-producers suggests that tuning the expression of this gene could potentially be a useful strategy for controlling EPO production in HEK293F cells.

In conclusion, the present study offers important insights into transcriptomic changes during secretory EPO and non-secretory GFP protein production as well as key parameters influencing the different rates of protein production across a variety of protein producers. The results are thus valuable for improved understanding of the biology behind protein secretion in mammalian cells and also the behaviour of cells under ER stress conditions. Moreover, the key differences uncovered between high- and low-producing EPO clones are potentially useful for future targeted cell-line engineering to improve therapeutic protein production.

## Acknowledgements

This work was supported by the Knut and Alice Wallenberg Foundation, AstraZeneca, Swedish Foundation for Strategic Research (SSF), Swedish innovation agency Vinnova through AAVNova, CellNova and AdBIOPRO and the Novo Nordisk Foundation (grant no. NNF10CC1016517).

## Author Contributions

Conceptualization, JN, JLR, and JR; Methodology, RS, JZ, MM, JLR and JR; Formal analysis, RS, JZ and MM; Investigation, RS, MM, JZ, VC, RR, NW, JLR and JR; Writing – Original Draft, RS, JZ, MM, JLR and JR; Writing – Review & Editing, RS, JZ, MM, DH, JN, LG, VC, JLR and JR; Visualization, RS, JZ, LG and JLR; Supervision, DH, TS, JN, JLR and JR, Funding Acquisition, JN and JR.

## Declaration of Interests

The authors declare no competing interests.

## STAR Methods

### Key Resources Table

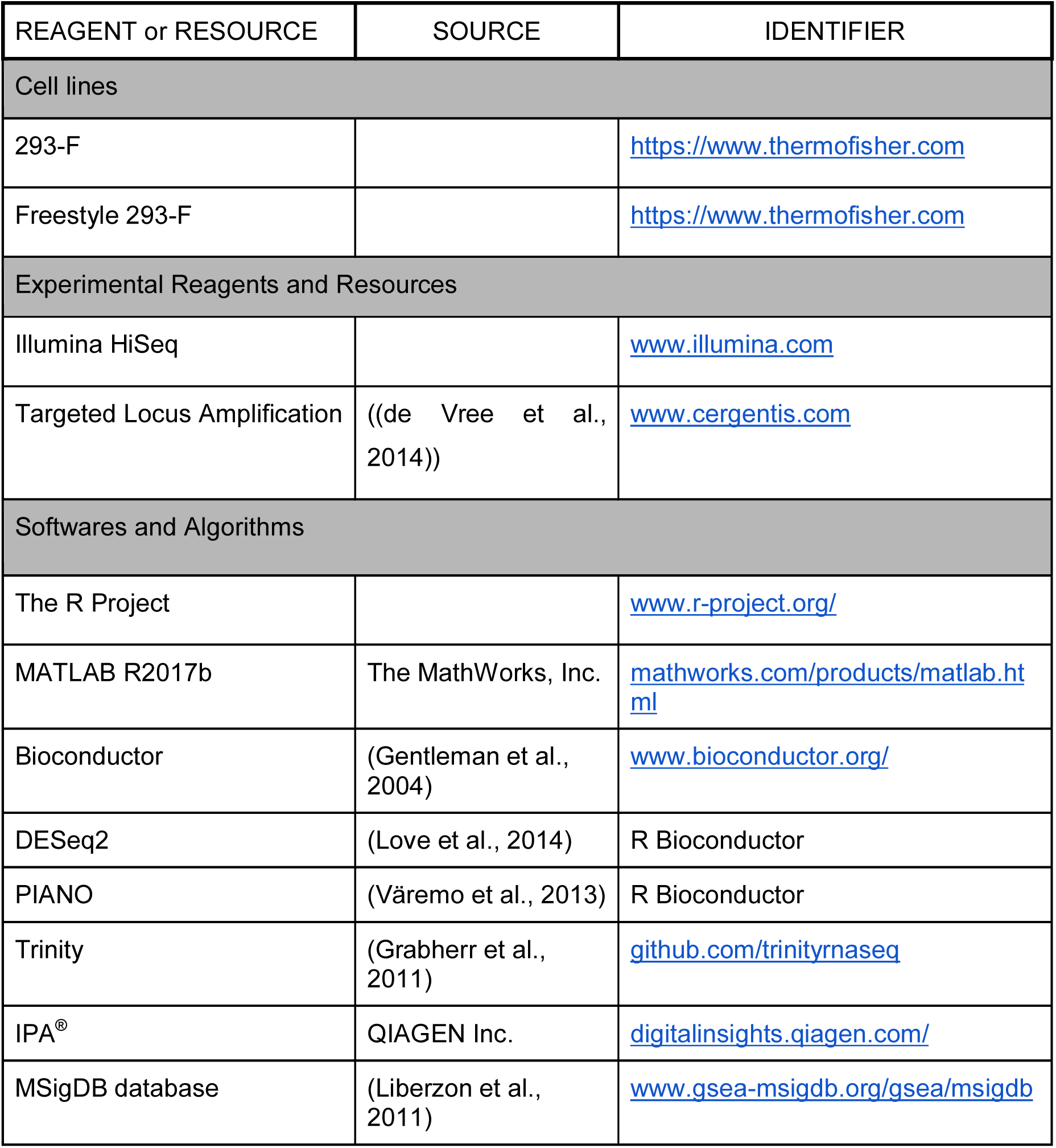

#### Contact for Reagent and Resource Sharing

Further information and requests for resources should be directed to and will be fulfilled by the Lead Contact, Johan Rockberg.

## Method Details

### M1. Cell cultivation and isolation of single clones

HEK293 cell lines 293-F and Freestyle 293-F were cultivated in Freestyle 293 expression medium (Gibco, Thermo Fisher Scientific) at 37°C, 125 rpm and 8% CO_2_. Stable cell clones expressing EPO or GFP were generated by transfection of linearized pD2529 plasmids (Atum), expressing either recombinant human EPO fused to a C-terminal HPC4-tag or recombinant GFP, into Freestyle 293-F (for EPO clones) or 293-F (for GFP clones) cell lines. Transfections were carried out using PEI at a DNA:PEI ratio of 1:3 and 1 μg plasmid per 1 million cells. Polyclonal batches of cells expressing GFP and EPO, respectively, were generated by puromycin selection. Single clones of HEK293 cells expressing EPO or GFP were generated from the polyclonal batches by seeding single cells per well of 384-well plates by either limited dilution or FACS (Astrios, Beckman Coulter). In case of GFP-expressing cells, sorting by FACS was performed based on the GFP-signal. Verification of cell monoclonality was performed by microscopy (Leica DMI6000B). Single cells were expanded in 1.5% HEPES and growth media at 37°C and 8% CO_2_. Cells of single clones were seeded at 0.3 million cells/ml in duplicates per clone and cultivated for 72 hours in 125 ml Erlenmeyer shake flasks with vented caps at 125 rpm, 37°C and 5% CO_2_. Every 24 hours cell growth and viability of cells were determined using a TC20 cell counter (Bio-Rad Laboratories). At 72 hours post inoculation, cell and supernatant samples were collected for downstream analysis. Cell samples for downstream RNA isolation were stored in RNAlater™ stabilization solution (Invitrogen™).

### M2. Productivity measurements

Specific productivity of EPO in cell supernatants was determined by Octet RED96 biolayer interferometry (ForteBio, Fremont, CA, USA) as described by (Kol et al., 2015). Briefly, biotinylated V_H_H-anti EPO (Capture Select™, Thermo Scientific) was immobilized on streptavidin sensors and used to measure EPO binding directly in cell supernatants in citric acid (20 mM), 0.1% BSA, 0.1% tween-20, 0.5 M NaCl. Signals were compared to an EPO standard curve of known concentration. Regeneration of sensors was performed using 10 mM NaH_2_PO_4_ (pH 12). GFP productivity was determined by measuring the GFP signal (FL-1 channel) of cells by flow cytometry (Gallios, Beckman Coulter).

### M3. Genome copy number estimation of GFP and EPO using TLA technology

Cryopreserved cell stocks, in cell growth medium with 10% DMSO, of each cell clone were sent to Cergentis B.V. (Utrecht, The Netherlands) for Targeted Locus Amplification (TLA) (de Vree et al., 2014) and next-generation sequencing by Illumina MiniSeq. EPO and GFP transgene sequences were mapped to the pD2529 plasmid sequences used for generating stable clones. Target-specific sequences were mapped to the hg19 genome (Genome Reference Consortium Human Build 37 (GRCh37). Estimations of copy numbers were based on number of plasmid integrations into the genome, number of fusion reads and ratio between coverage of transgene and surrounding genomic region.

### M4. Correlation analysis

For correlation analyses between growth rates, protein productivity, gene and transcript copy number and final visualizations, the R package GGally v1.5.0 (https://github.com/ggobi/ggally) was used with default settings. Correlation analysis between gene expression and EPO or GFP production was performed using the Pearson correlation coefficient, where results were filtered based on the significance threshold of 0.05 and absolute correlation coefficients surpassing 0.5. An additional filter was applied to exclude genes with a mean of TPM lower than 10 across all samples of the same producing group. Interaction between correlating genes was based on the STRING database (Szklarczyk et al., 2019) and filtered according to experimental evidence and a confidence interval higher than 900. First-order interacting partners of correlating genes with mean of TPM higher than 10 also included in the networks. For network visualization and gene set analysis of highly correlated genes, NetworkAnalyst (Zhou et al., 2019) was used with default settings and GO biological process version v7.1 was used as the gene set collection.

### M5. RNA preparation and sequencing and data analysis

Total RNA was extracted from cells using RNeasy plus Mini Kit (Qiagen) according to the protocol provided by the manufacturer. RNA integrity was verified by RNA 6000 Nano chips on a 2100 Bioanalyzer instrument (Agilent Technologies). Extracted RNA samples were shipped to GATC (Konstanz, Germany) for mRNA sequencing by Illumina HiSeq instrument using the Inview Transcriptome Discover service (paired end, 2 × 150 bp read length, >30 million read pairs).

Transcript quantification was performed using the standalone package Kallisto v0.43.1_1 (Bray et al., 2016) with default settings and version GrCh38 of the human genome (Schneider et al., 2017) was used for transcript mapping. To import raw counts data into R, the tximport package v1.14.2 (Soneson et al., 2015). For PCA analysis, the R package DESeq2 v1.26.0 (Love et al., 2014) was used with default settings and log transformed normalized counts. Differential expression analysis was also performed using DESeq2 v1.26.0, where the Wald test was used for calculating logarithmic fold changes and *p*-values were adjusted by the Benjamini-Hochberg method. To find genes with a robust expression change between EPOF21 and other EPO producer clones, we filtered genes with an absolute log2 fold change higher than 1, adjusted *p*-value smaller than 0.05 and average TPM higher than 10 across all samples.

### M6. Ingenuity pathway analysis

Enrichment analysis for finding enriched pathways was performed using Ingenuity Pathway Analysis (IPA^®^) software (Ingenuity Systems, http://ingenuity.com) with default settings. For all comparisons, gene names were first mapped to the Ingenuity database and then statistically significant differentially expressed genes (B.H. adj. *p*-value < 0.05) with at least 50% change in their expression were selected for finding significantly enriched pathways (Benjamini-Hochberg corrected *p*-value < 0.05).

### M7. Gene set Analysis

To measure the enrichment of different gene groups, gene set analysis (GSA) was performed. The procedure for running GSA was the same across the study. First, gene set collections were retrieved from the Molecular Signature Database (MSigDB) (Liberzon et al., 2011). For gene sets related to Human1 (Robinson et al., 2020), the MATLAB toolbox RAVEN v2.3.1 was used (Wang et al., 2018) with default settings to group genes based on their associations with reactions in different subsystems of the Human1 v1.3.1 model. To calculate test statistics for each given gene set, the Wilcoxon rank-sum from the R package PIANO v2.2.0 (Väremo et al., 2013) was used, where results for each gene set based on *p*-values and log2 fold-change of genes from the DE analysis were compared with 100,000 randomly shuffled gene sets of the same size (random permutations = 100,000).

### M8. Generating GO slim secretion

To estimate the extent to which the protein secretory subsystems differed between EPOF21 and other EPO producers, a GO slim for protein secretion-related gene subsets was generated for use in gene set analysis. The GO slim consisted of a list of 590 previously reported genes involved in human protein secretion and their association to core components of the secretory pathway (Feizi et al., 2017). In addition, gene sets were retrieved from the GO biological processes, GO cellular components and GO molecular functions MSigDB collections (Liberzon et al., 2011) if they were significantly enriched in secretion-associated genes (B.H. adj. *p*-value < 0.05). A one-tailed Fisher’s exact from the R package PIANO v2.2.0 was used as the statistical test for calculating the enrichment significance of gene sets and *p*-values were adjusted to control for the false discovery rate (*FDR*) with the Benjamini-Hochberg procedure.

### M9. Software

All computational analyses were performed using R v3.6 (www.r-project.org) and Matlab R2017b (www.mathworks.com). Code and datasets to reproduce the figures presented here as well as all analysis outputs, are available on GitHub repository: https://github.com/SysBioChalmers/EPO_GFP. Data files too large to host on GitHub were deposited on Zenodo: https://zenodo.org/record/4004264#.X0iqstMzbuM

## Supplemental Information titles and legends

**Figure S1.**
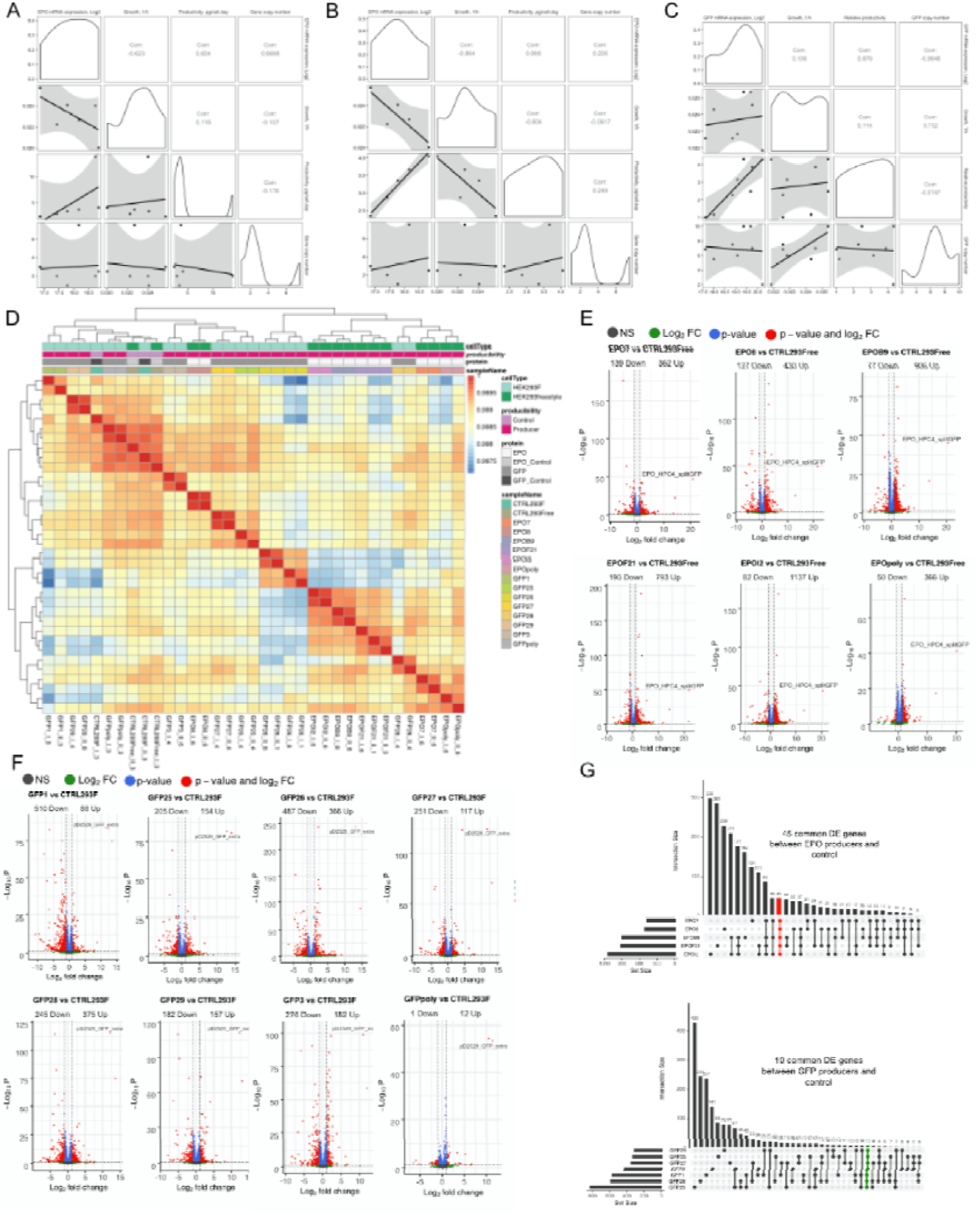
Correlation analysis between growth rate, protein productivity, gene and mRNA copy number. (A) EPO producer clones and (B) EPO producers without the highest producing EPOF21 clone. (C) Correlation analysis between growth rate, protein productivity, gene and mRNA copy number for GFP producer clones. (D) Samples correlation plot for all replicates. (E) Differentially expressed genes (B.H. adj. *p*-value < 0.05, | L2FC | > 1) between EPO producers vd. control (E) and GFP producers vs control (F). (G) Common differentially expressed genes (*p*-value < 0.05, | L2FC | > 1) in pairwise comparison of EPO producer clones wirth control (45 genes) and pairwise comparison of GFP producer clones wirth control (10 genes).

**Figure S2.**
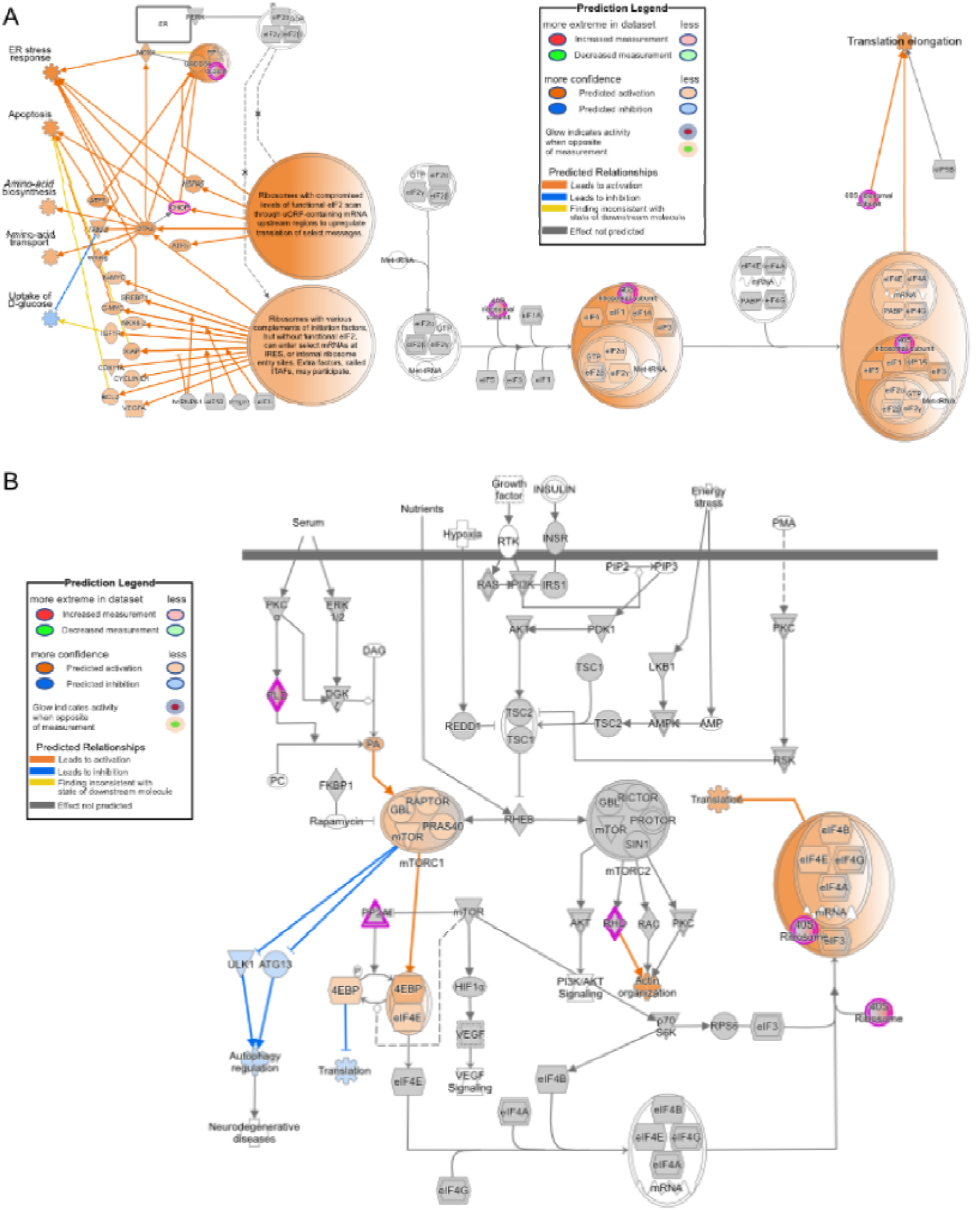
eIF2 signaling and mTOR signaling pathways show activation of translation in EPO producers in comparison to host parental cell-line. Detailed visualization of mTOR signaling and Eukaryotic Initiation Factor 2 (eIF2) signaling pathways that were the most significantly differentially up regulated pathways across all EPO producer clones compared to control cell-line.

**Figure S3.**
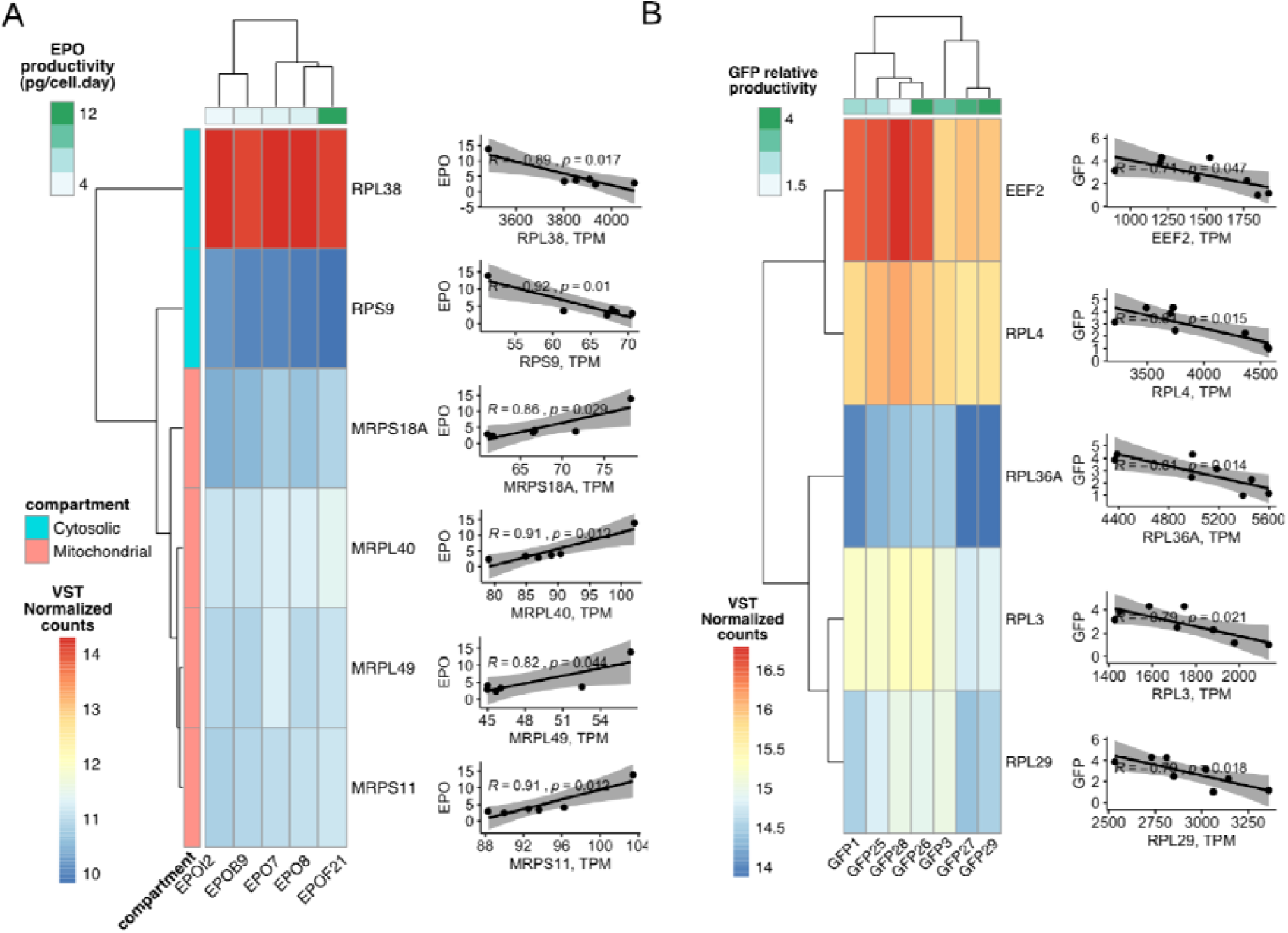
Cytosolic and mitochondrial ribosomal genes have a reverse trend of expression in EPO producers. (A) RPL38 and RPS9 ribosomal genes have significant negative correlation with EPO production and both are components of cytosolic ribosomes while MRPS18A, MRPL40, MRPL49 and MRPS11 are components of mitochondrial ribosomes and all have significant positive correlation with EPO production. (B) All ribosomal genes with significant correlation with GFP are components of cytosolic ribosome and all have negative correlation with GFP production.

**Figure S4.**
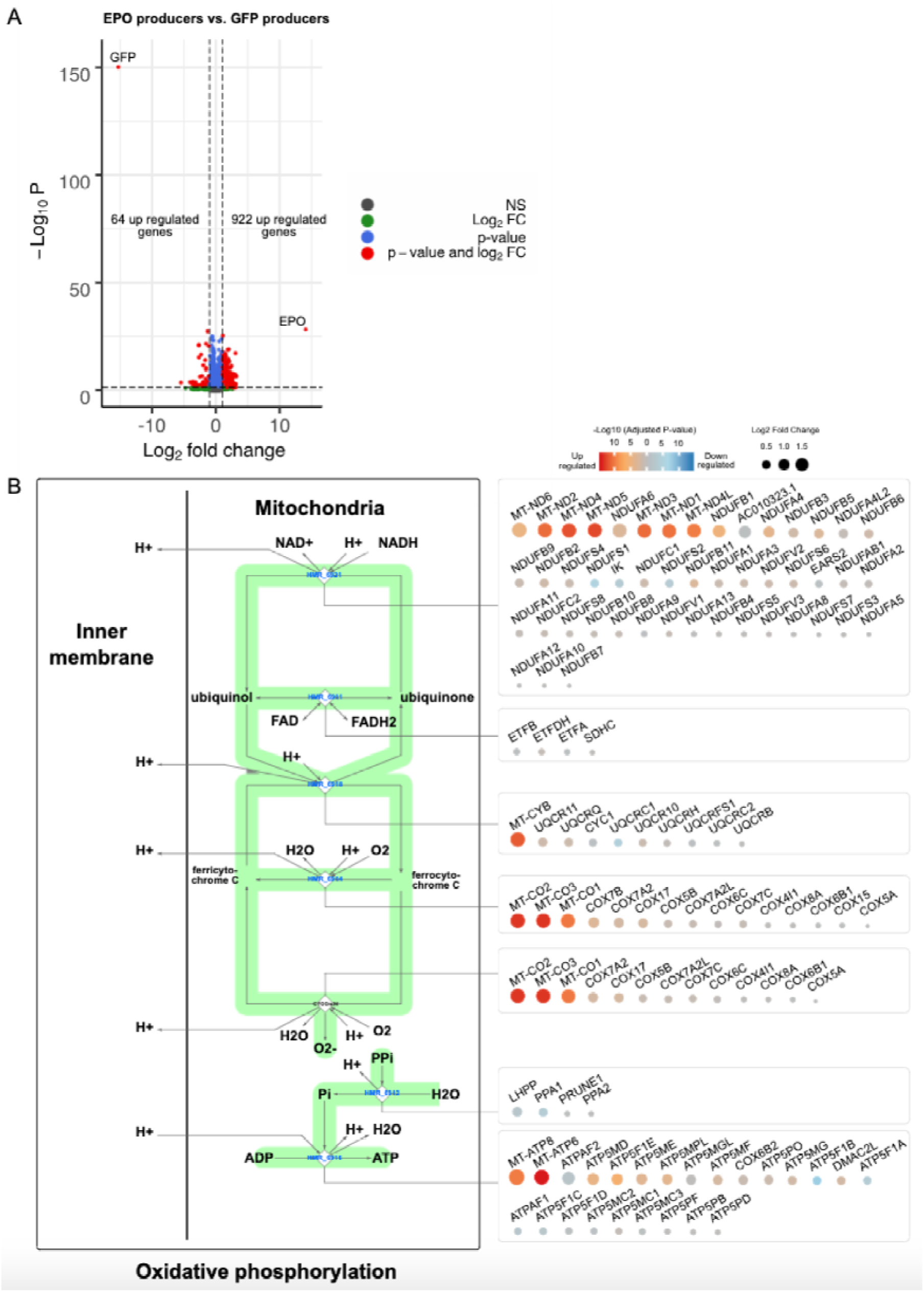
Increase in energy production in EPO clones is governed by the upstream regulatory pathways of the cell. Apart from up regulation of mitochondrial encoded genes, other genes like NDUFA6, NDUFB1 and NDUFA4 in complex I, COX7B and COX7A2 in complex III and ATP5MD and ATP5F1E in complex V have a significant (B.H. adj. *p*-value < 0.05) expression increase in EPO producers compared to GFP producers.

**Figure S5.**
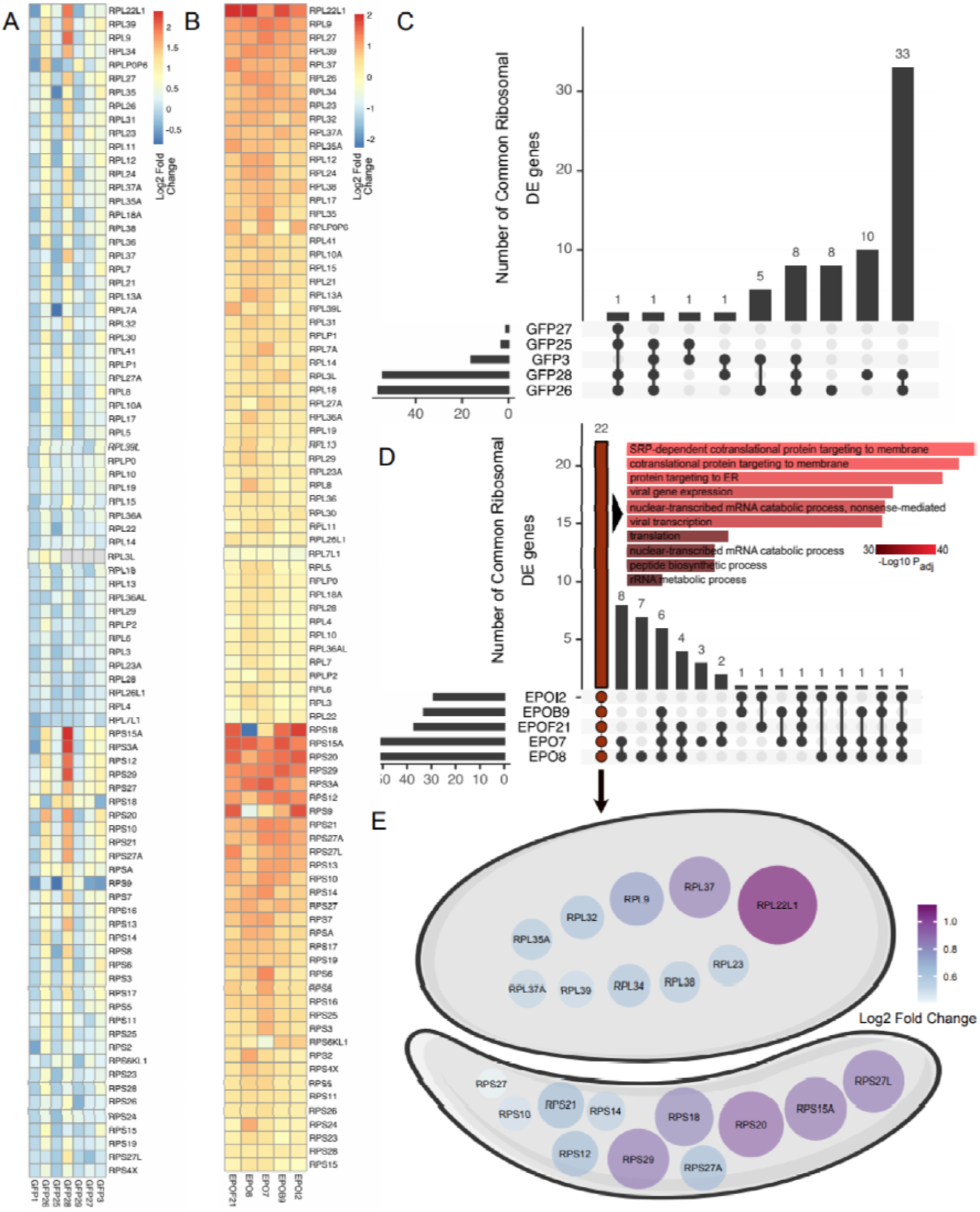
Ribosomal genes show recombinant protein specific patterns of expression. (A) Pairwise comparison of ribosomal genes between GFP producers versus control. (B) Pairwise comparison of ribosomal genes between EPO producers versus control. (C) Common significantly differentially expressed (B.H. adj. *p*-value < 0.05, | L2FC | > 0.58) ribosomal genes across all GFP producers compared to control. (D) 22 common significantly differentially expressed (B.H. adj. *p*-value < 0.05, | L2FC | > 0.58) ribosomal genes across all EPO producers in comparison to control. Most significantly associated pathways with these genes belong to SRP (signal recognition particle) dependent cotranslational protein targeting to the membrane. (E) Fold change of common differentially expressed ribosomal genes between EPO producers versus control.

**Figure S6.**
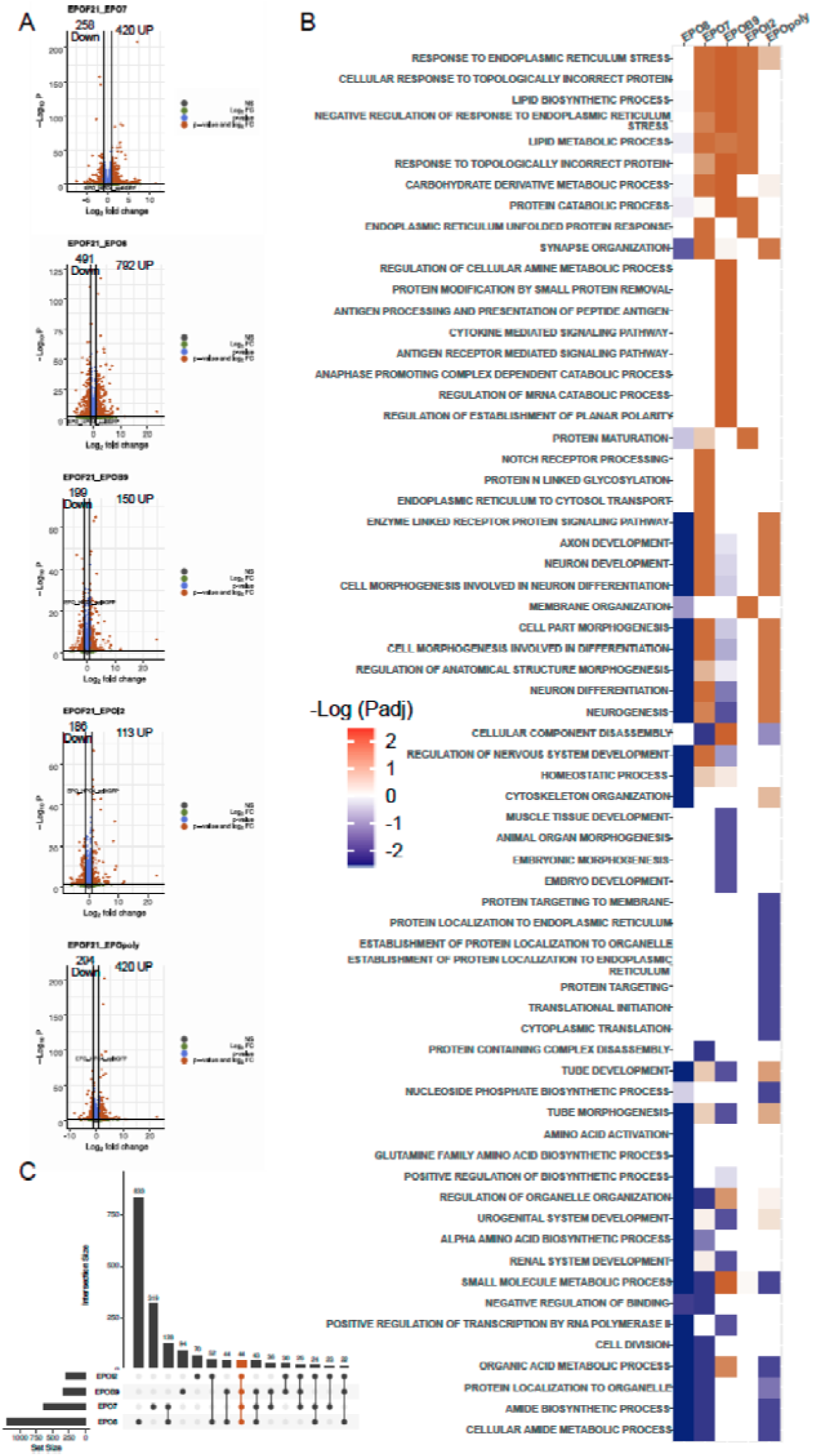
Gene enrichment analysis between EPOF21 and other low EPO producers highlights differences in post-translational pathways. (A) Differentially expressed genes between EPO producers vs. EPOF21 high producer clone. EPO expression is not highly changed between EPO producers (B) Enriched pathways for differentially expressed genes in pairwise comparison of EPOF21 with each of low EPO producer clones. (C) There are 44 common differentially expressed genes between EPOF21 and other EPO producers.

**Figure S7.**
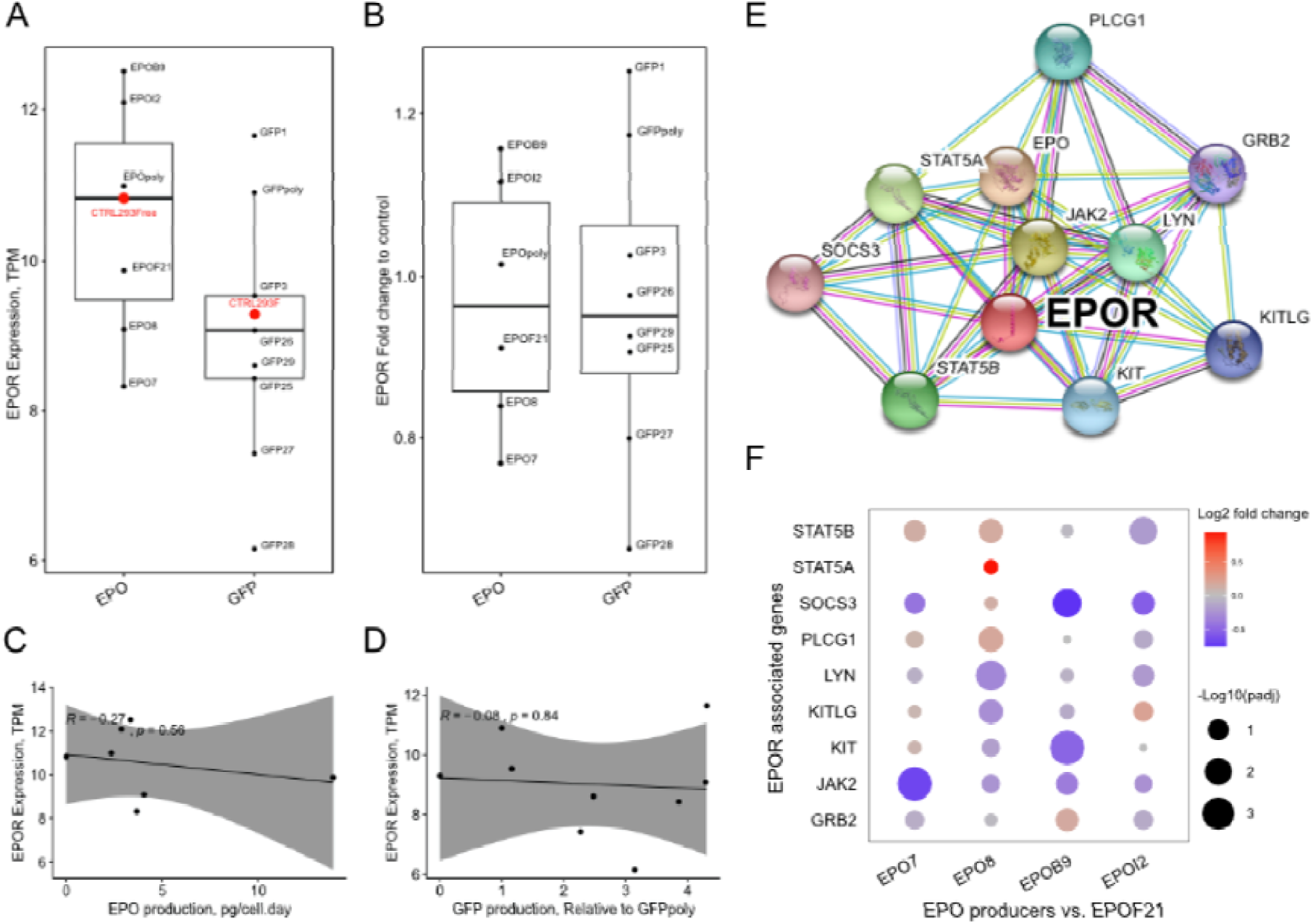
The difference in expression of Erythropoietin receptor (EPOR) gene is not significant between EPO producers vs. control cell-line and also EPO producers vs. GFP producers. (A) Distribution of EPOR expression between EPO producers and GFP producers. (B) Normalized expression of EPOR by considering control cell-lines background effect shows no difference in expression of EPOR between EPO producers and GFP producers. (C) EPOR expression is not significantly correlated with EPO production across EPO producer clones including control cell-line (D) There is no significant correlation in EPOR expression and GFP production in GFP producer clones. (E) Network of interacting genes with EPOR. (F) Comparison of expression of EPOR interacting genes indicates expression of these genes is not significantly different (B.H adj. *p*-value < 0.05, L2FC > 1) between EPOF21 and other EPO producer clones.

**Table S1: Table of raw counts for transcriptomics analysis**

**Table S2: List of enriched pathways in pairwise comparison of EPO producers against EPO control host.**

**Table S3: List of significantly correlated genes with EPO and GFP production and statistics for EPO- and GFP-correlating genes networks**

## Notes

https://github.com/SysBioChalmers/EPO_GFP

https://zenodo.org/record/4004264#.X0iqstMzbuM

### Competing Interest Statement

The authors have declared no competing interest.

