## Supplementary figures and images for "Transcriptome analysis of EPO and GFP HEK293 Cell-lines Reveal Shifts in Energy and ER Capacity Support Improved Erythropoietin Production in HEK293F Cells"

### Supplemental Figure 1

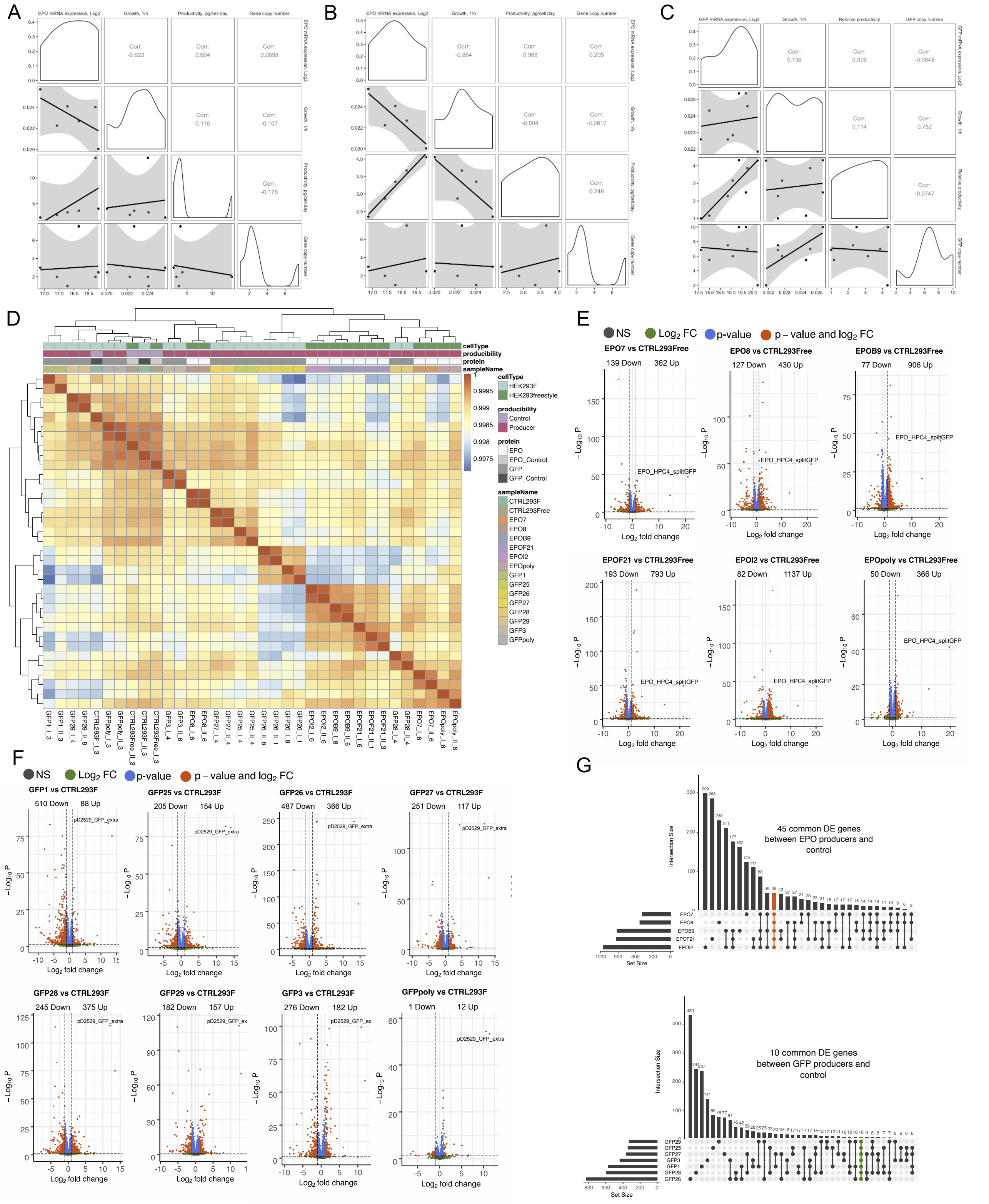

### Supplemental Figure 2

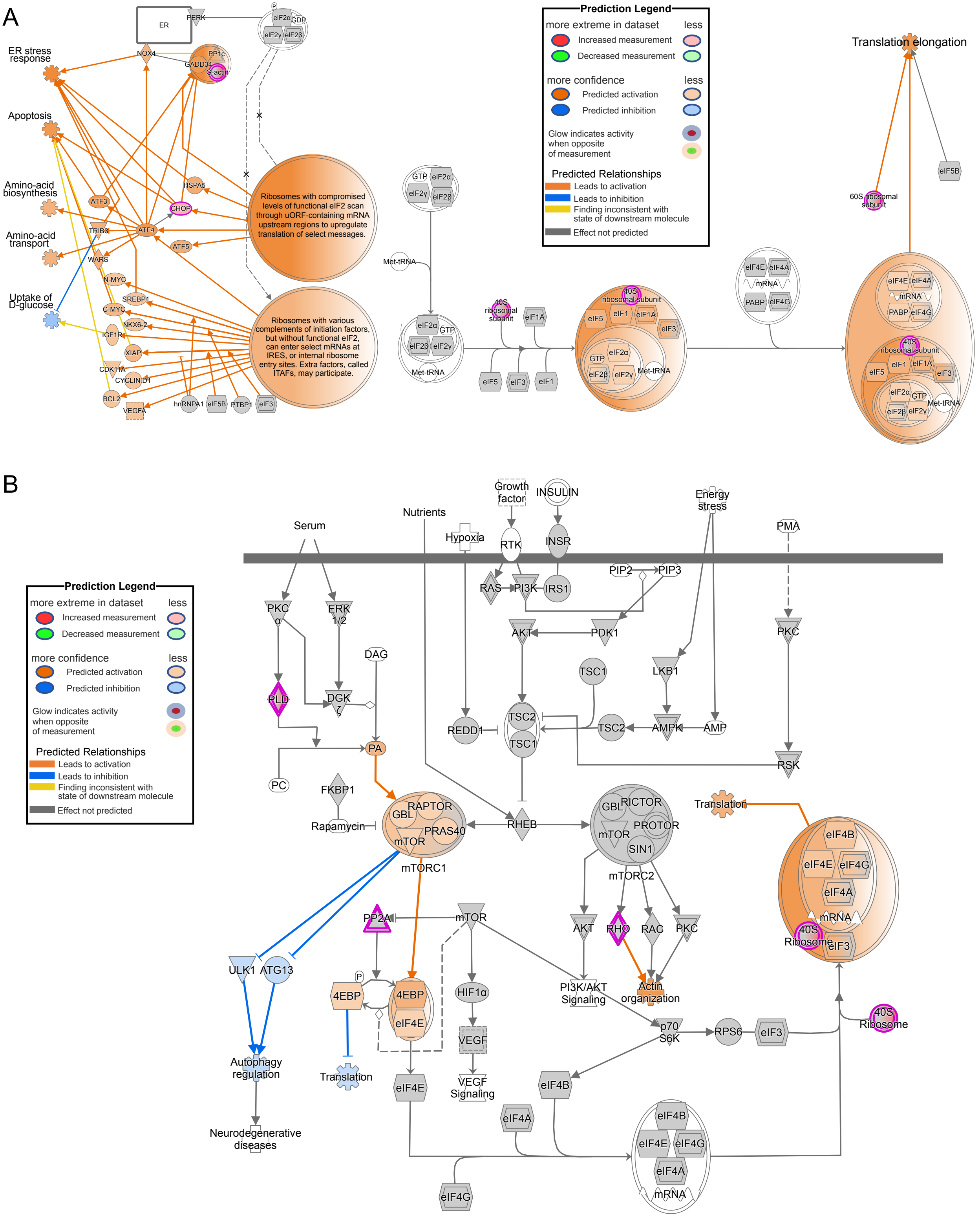

### Supplemental Figure 3

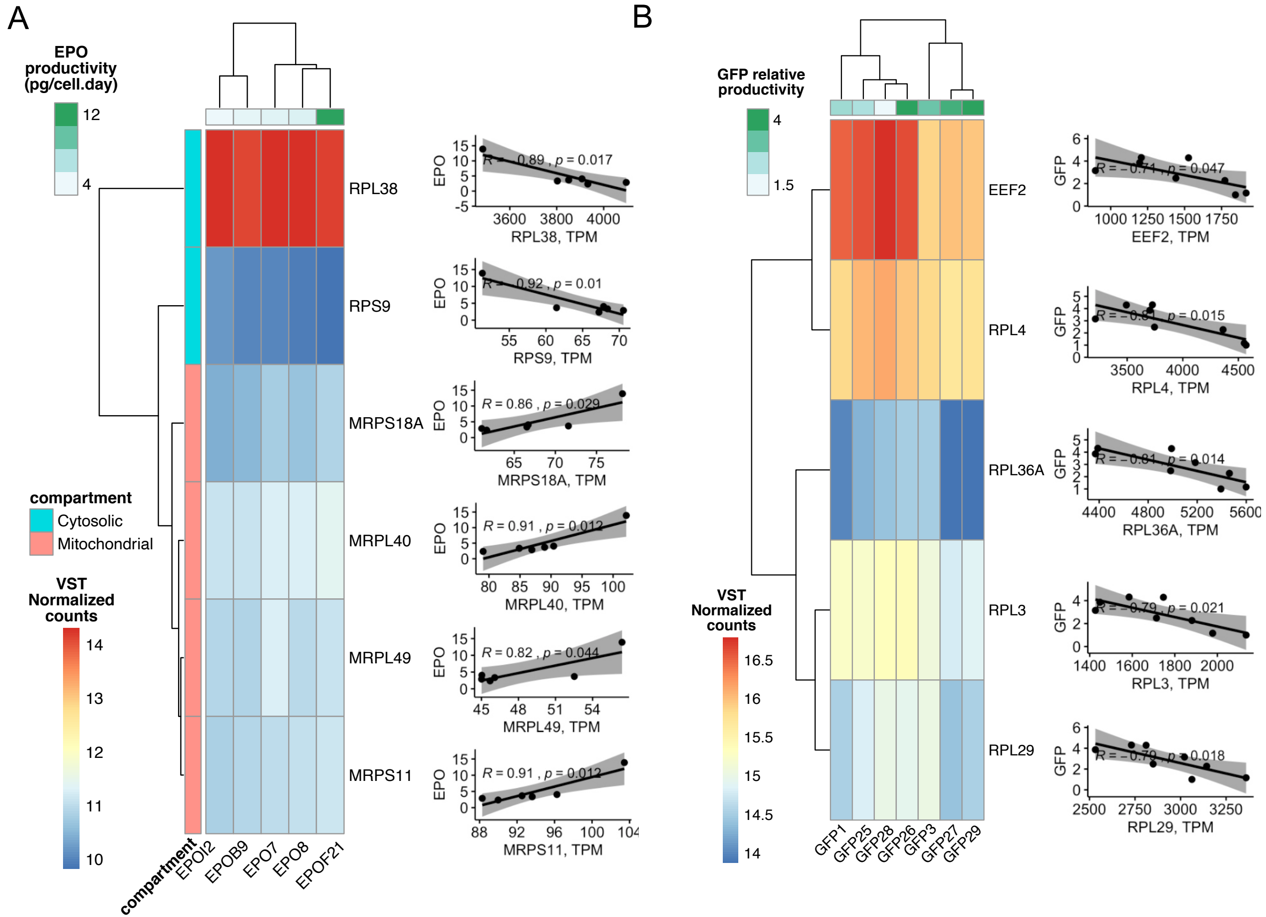

### Supplemental Figure 4

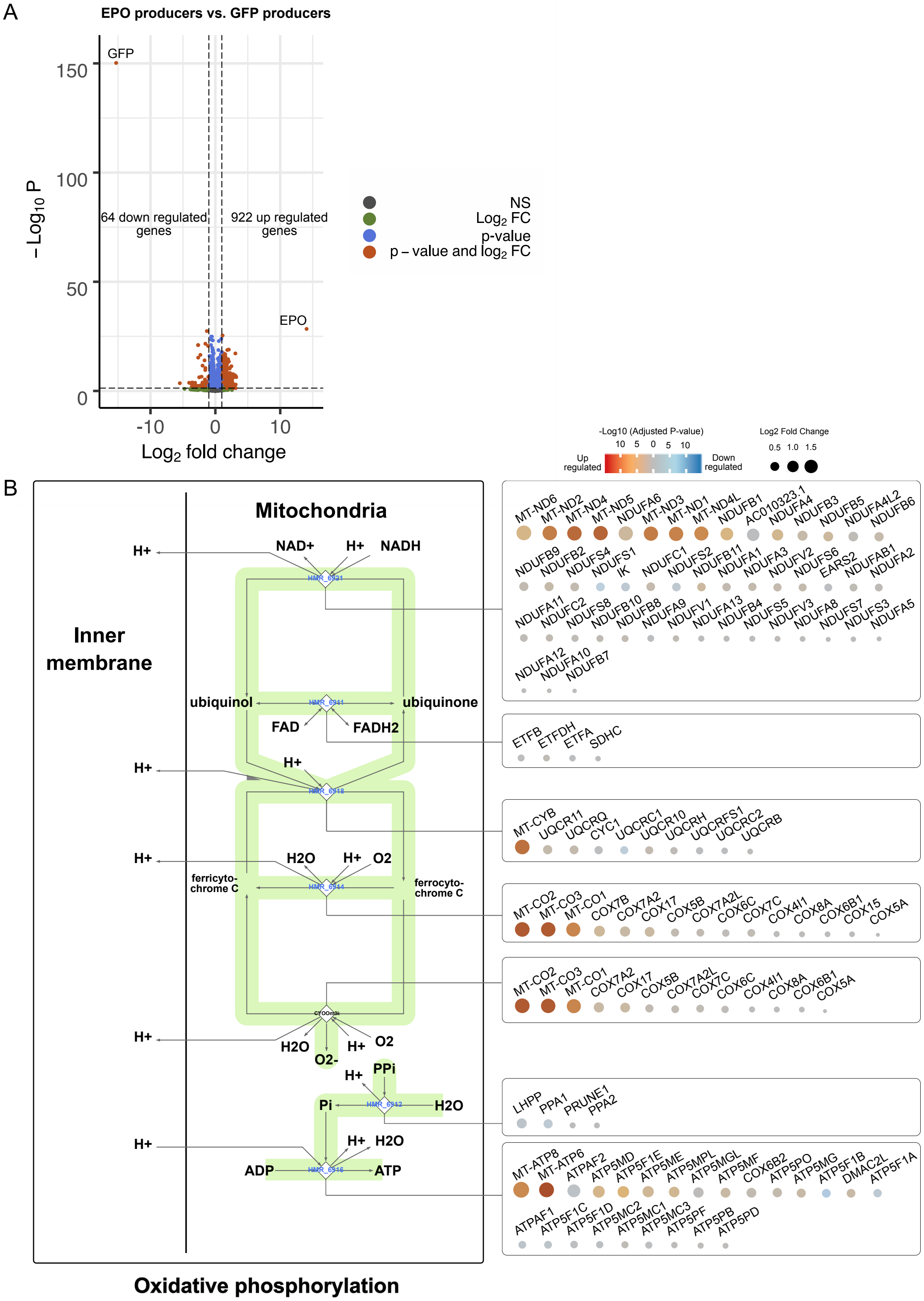

### Supplemental Figure 5

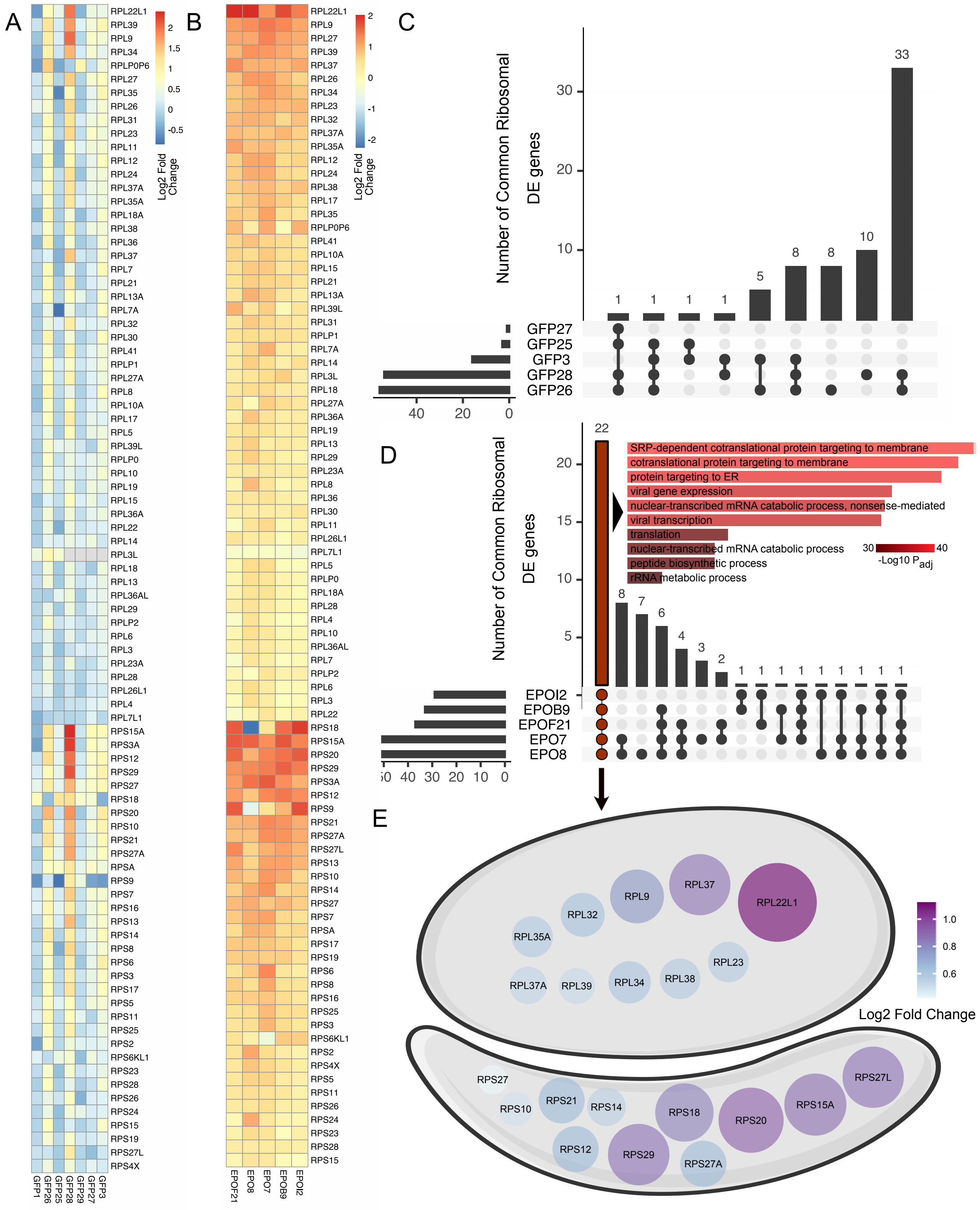

### Supplemental Figure 6

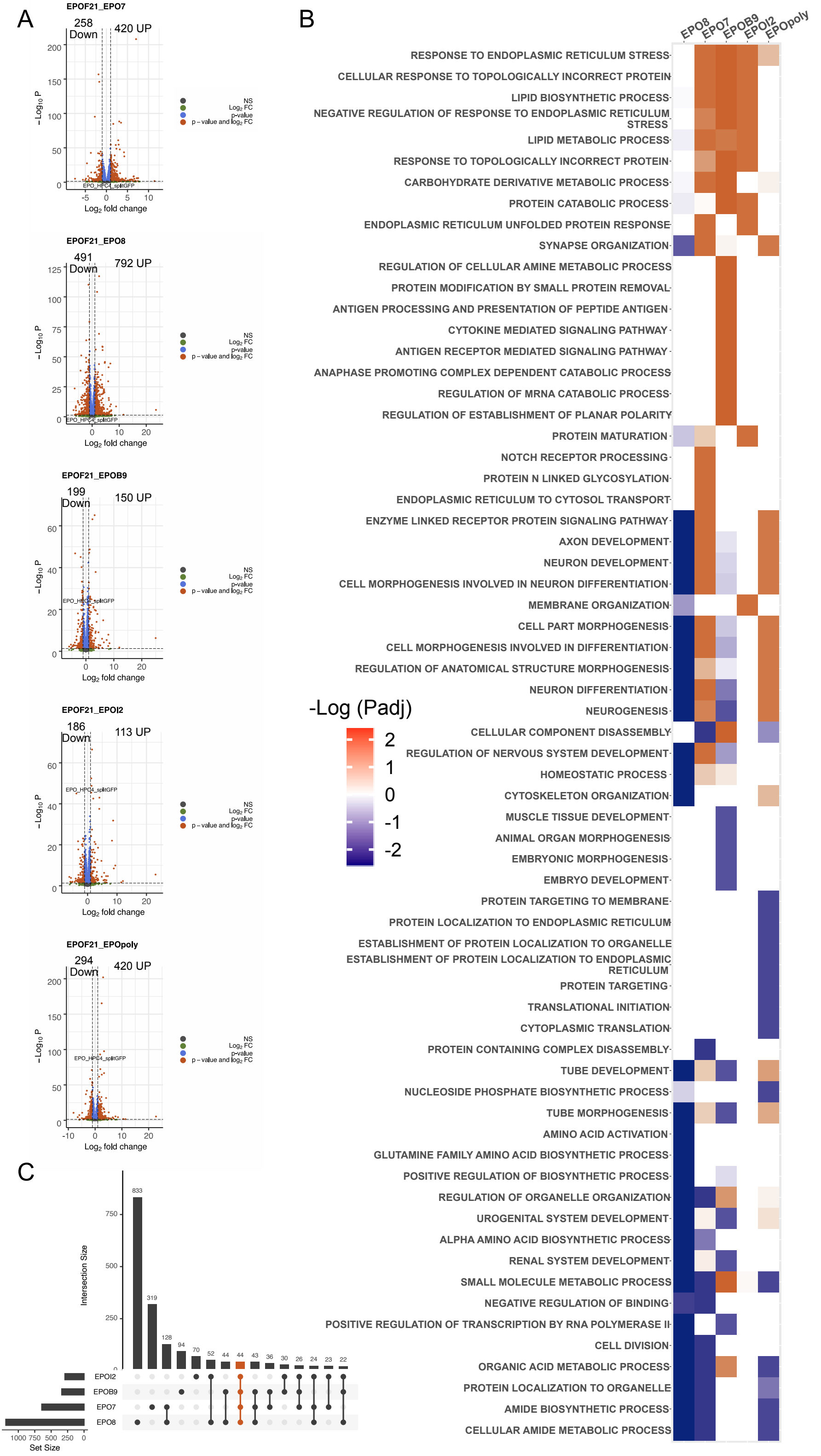

### Supplemental Figure 7

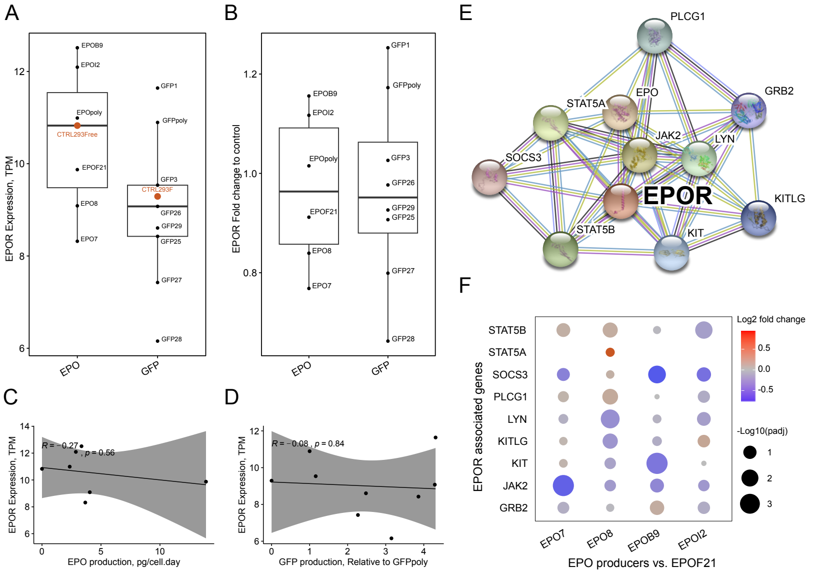
